# Modelling human endurance: Power laws vs critical power

**DOI:** 10.1101/2022.08.31.506028

**Authors:** Jonah Drake, Axel Finke, Richard Ferguson

## Abstract

The *power–duration relationship* describes the time to exhaustion for exercise at different intensities. It is generally believed to be a “fundamental bioenergetic property of living systems” that this relationship is hyperbolic. Indeed, the *hyperbolic* (a.k.a. *critical-power*) model which formalises this belief is the dominant tool for describing and predicting high-intensity exercise performance, e.g. in cycling, running, rowing, or swimming. However, the hyperbolic model is now the focus of two heated debates in the literature because: (a) it unrealistically represents efforts that are short (< 2 minutes) or long (> 15 minutes); (b) it contradicts widely-used performance predictors such as the so-called *functional threshold power* (*FTP*) in cycling. We contribute to both debates by demonstrating that the power–duration relationship is more adequately represented by an alternative, *power-law* model. In particular, we show that the often observed good fit of the hyperbolic model between 2 and 15 minutes should not be taken as proof that the power–duration relationship is hyperbolic. Rather, in this range, a hyperbolic function just happens to approximate a power law fairly well. We also prove mathematical results which suggest that the power-law model is a safer tool for pace selection than the hyperbolic model and that the former better models fatigue than the latter. Finally, we use the power-law model to shed light on popular performance predictors in cycling, running and rowing such as FTP and Jack Daniels’ *“VDOT” calculator*.

## 1 Introduction

### 1.1 The power–duration relationship

The *power–duration relationship* describes the time to exhaustion for exercise at different intensities. Knowledge of this relationship is crucial to athletes and coaches, e.g. for:

- *fitness assessment* – i.e. in order to inform training, athletes (and their coaches) want to quantify and track their fitness level;
- *performance prediction* – i.e. in order to select race and pace strategies, athletes want to predict their potential over distances or durations that may not have recently been performed. For instance,
  – runners want to know their best possible finish time in a marathon from only a recent half-marathon performance (i.e. without running the full distance);
  – cyclists want to know if they can sustain a particular power output for a given distance or duration (e.g. needed at the final climb of a race).

In this work, we focus on the endurance sports: running, cycling and rowing, although we stress that our results are not limited to these activities and may even be relevant in other settings, e.g. functional capacity testing in clinical populations.

Unfortunately, an individual’s power–duration relationship is unknown and must be estimated from available data with the help of mathematical models that formalise the power–duration relationship. A number of such models have been proposed in the literature. Among these are the following two which were both originally used to model world-record performances *across* different athletes (and even animals) but which are nowadays also used to describe the power–duration relationship of individual athletes.

- **Hyperbolic (a.k.a. critical-power) model.** The *hyperbolic* model (Hill, 1925; Monod and Scherrer, 1965; Hill, 1993; Jones et al., 2010) – illustrated in Figure 1 – asserts that the power–duration relationship is hyperbolic^1^. The power asymptote is called *critical power*. The model is thus also called the *critical-power model*.
- **Power-law (a.k.a. Riegel) model.** The *power-law* model (Kennelly, 1906) – illustrated in Figure 2 – asserts that the power–duration relationship follows a power law. It was popularised in running by Riegel (1977, 1981). The model is thus often termed the *Riegel* model.

**Figure 1:**
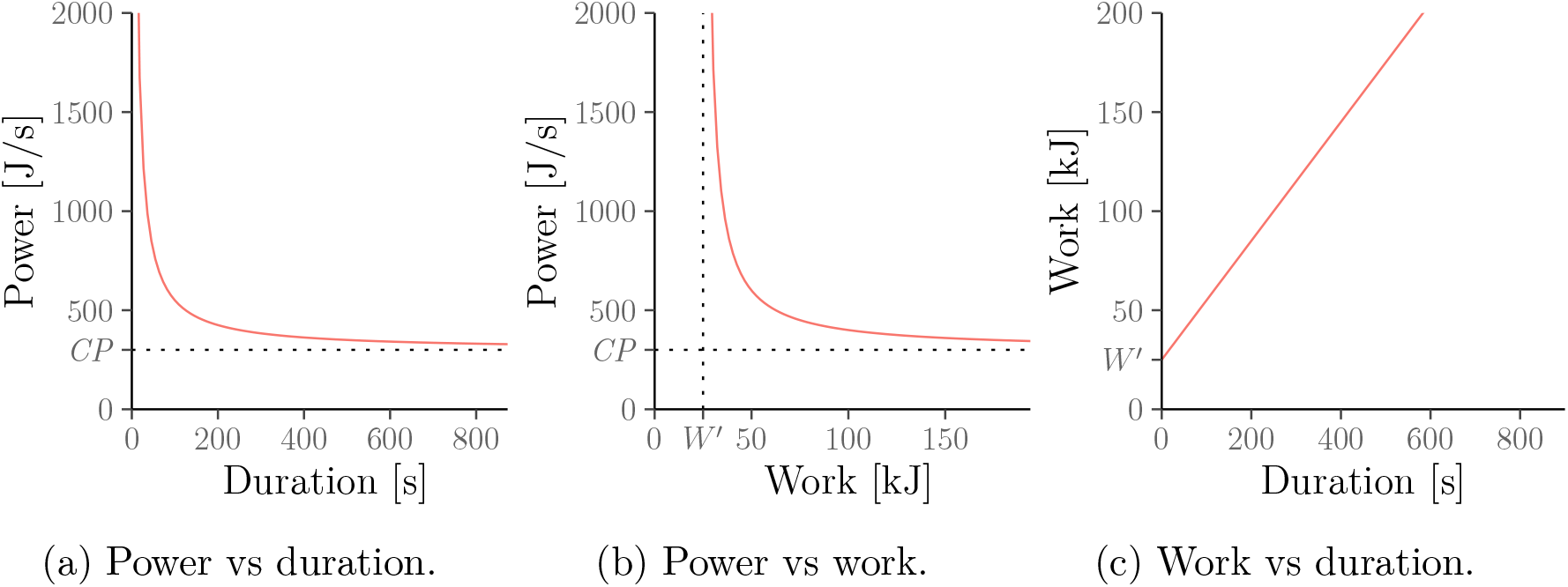
Equivalent relationships between power, duration and work under the hyperbolic (a.k.a. critical-power) model. Note that the power output approaches the critical power CP > 0 in Panels a and b as duration and work increase.

**Figure 2:**
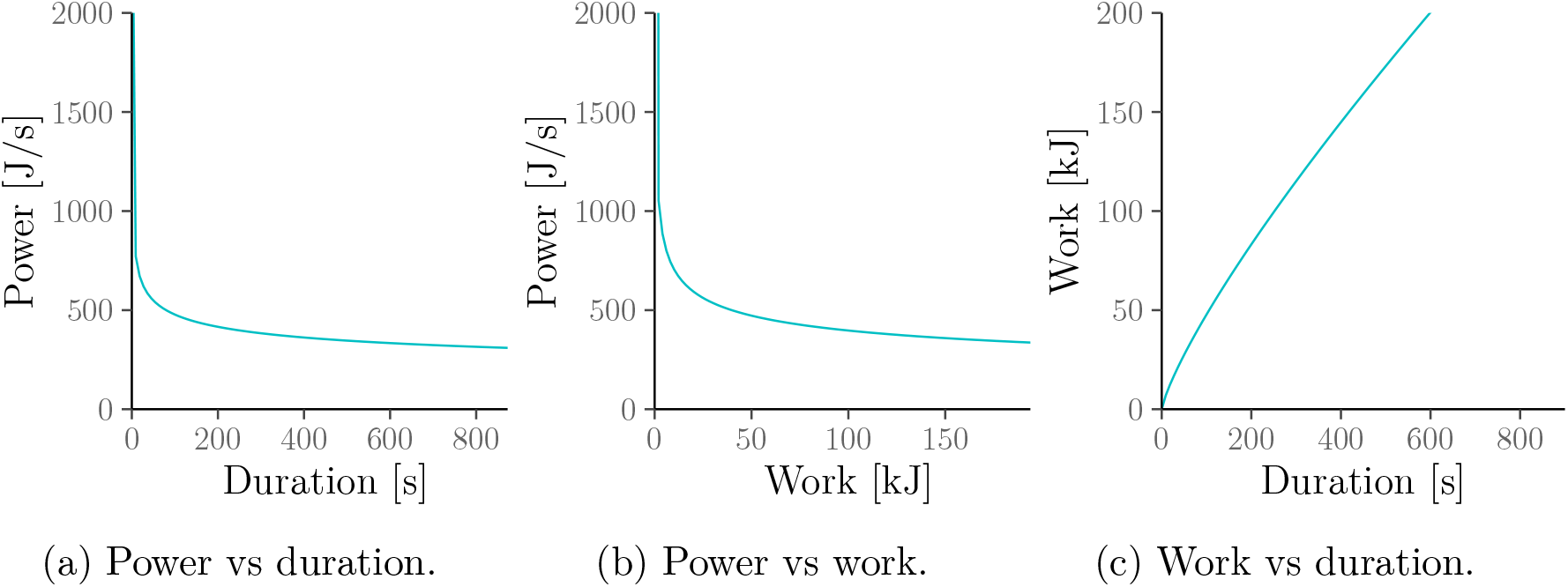
Equivalent relationships between power, duration and work under the powerlaw (a.k.a. Riegel) model. Note that the power output approaches 0 in Panels a and b as duration and work increase. This is in contrast to the hyperbolic (a.k.a. critical-power) model in Figure 1 where it approaches CP > 0.

### 1.2 Recent debates about the critical-power paradigm

The hyperbolic (a.k.a. critical-power) model is considered to describe and predict endurance performance during high-intensity exercise (i.e. exercise classed as “severe”) “with startling precision” (Poole et al., 2016). It is ubiquitous in cycling (Leo et al., 2022a), where it is implemented in popular online exercise analytics platforms; but it is also widely used for training prescription and performance prediction in other endurance sports such as running (Kranenburg and Smith, 1996; Nimmerichter et al., 2017), rowing (Hill et al., 2003), swimming (Wakayoshi et al., 1992; Petrigna et al., 2022) as well as walking and skating (Hill, 1925). Additionally, it has been applied to intermittent sports such as football, hockey and rugby (Okuno et al., 2011) in order to optimise the length of recovery needed between exercise bouts (Fukuda et al., 2011). The model has also been suggested as a tool for anti-doping (Puchowicz et al., 2018) and for assessing the effectiveness of nutritional supplements (Fukuda et al., 2010; Stout et al., 1999).

The hyperbolic model has even been employed to measure exercise capacity in horses and mice (Lauderdale and Hinchcliff, 1999; Billat et al., 2005). Indeed, the hyperbolic shape of the power–duration relationship which the model formalises is considered to be a “fundamental bioenergetic property of living systems” (Jones et al., 2019).

However, the hyperbolic model is now at the centre of two controversies:

- **Unrealistic behaviour over short and long durations.** The first debate surrounds criticism that the hyperbolic model does not adequately describe the full power–duration relationship (Vandewalle et al., 1997) because the model behaves unrealistically for exercise durations that are short (e.g., less than 2 minutes) or long (e.g., more than 15 minutes). Indeed,
  – over short durations, the hyperbolic model would imply that an elite runner like Eliud Kipchoge can sustain over 650km/h over one second or instantaneously “teleport” over 175m;
  – over long durations, the hyperbolic model would imply that runners can complete ultra-marathons at essentially the same average velocity as half-marathons.
  This debate about the hyperbolic model has recently intensified with the publication of the papers by Dotan (2022a); Gorostiaga et al. (2022b) and the ensuing discussions (Burnley, 2022a,b; Black et al., 2022; Broxterman et al., 2022; Triska and Karsten, 2022; Marwood and Goulding, 2022; Altuna and Hopker, 2022; Lindinger, 2022; Dotan, 2022b,c; Gorostiaga et al., 2022c,a).
- **Incompatibility with FTP.** The second debate surrounds the fact that the hyperbolic model contradicts a metric known as *functional threshold power* (*FTP*) (Allen and Coggan, 2012) which is widely used in cycling (Morgan et al., 2019). Similarly, it disagrees with widely used rules-of-thumb for performance predictions such as Jack Daniels’ “*VDOT” calculator* (Daniels and Gilbert, 1979) and Jeff Galloway’s *Magic Mile* (Galloway, 1984) in running as well as Kurt Jensen’s *Golden Standard* (ergrowing.com, nd) in rowing.

### 1.3 Contributions

We show that the power–duration relationship should not be presumed to be hyperbolic because it is more adequately described by the power-law model; and that doing so resolves both the above-mentioned controversies about the critical-power paradigm.

Specifically, our novel contributions – both empirical and theoretical – are fourfold:

- **Contribution I: Fit over different durations.** In Section 3, we compare the fit of the power-law and hyperbolic models in the context of (a) the case studies of elite athletes from Jones and Vanhatalo (2017); Jones et al. (2019) and (b) additional large data sets of runners, rowers and cyclists, and demonstrate that:
  – for exercise durations *inside* the 2–15 minute range (for which the hyperbolic model is thought to be valid) the power-law model fits just as well as the hyperbolic model;
  – for exercise durations *outside* the 2–15 minute range (for which the hyperbolic model predicts performances that are physically impossible), the power-law model still exhibits realistic behaviour.
  This suggests that the good performance of the hyperbolic model in the 2–15 minute range cannot be taken as evidence for the hypothesis that the power–duration relationship is hyperbolic. Rather, the power–duration relationship is more adequately described by a power law and the hyperbolic model just happens to approximate the power-law model reasonably well in the 2–15 minute range.
- **Contribution II: Implications for pacing.** In Section 4, we prove a mathematical result which suggests that the power-law model has more realistic implications about pacing than the hyperbolic model. That is, the power-law model implies that athletes will only achieve their best possible finish time in a time trial if they race at the highest *constant* velocity that they can sustain until the finish line; and overpacing is always detrimental. In contrast, the hyperbolic model implies that the safest optimal pacing strategy is to sprint off as quickly as possible and then “hold on” until crossing the finish line. Thus, the power-law model is consistent with the pacing recommended by coaches and observed in elite athletes whereas the hyperbolic model is not.
- **Contribution III: Modelling fatigue.** In Section 5, we demonstrate that, unlike the hyperbolic model, the power-law model is automatically consistent with the empirical observation that the power–duration curve moves downwards as the athlete becomes fatigued from prolonged exercise. We also explain why the often-made assumption that critical power decreases with fatigue is contradictory.
- **Contribution IV: Relationship with popular performance predictors.** In Section 6, we show that various rules-of-thumb for performance prediction widely used by practitioners in cycling, running and rowing can be viewed as approximations of the power-law model, namely:
  1. *functional threshold power* (*FTP*) (Allen and Coggan, 2012);
  2. Jack Daniels’ “*VDOT” calculator* (Daniels and Gilbert, 1979);
  3. Jeff Galloway’s *Magic Mile* (Galloway, 1984);
  4. Kurt Jensen’s *Golden Standard* (ergrowing.com, nd).
  Our work therefore clarifies the strengths and limitations of these rules. That is, it formalises why these rules are likely biased for most athletes but why they can nonetheless be justified as a pragmatic *bias–variance trade-off* when few data points are available (meaning that in this case, it is worth accepting the bias because doing so reduce the overall prediction error).

### 1.4 Related literature

The finding that the power–duration relationship is more adequately modelled by a power law than a hyperbolic function has already been demonstrated in running (Hinckson and Hopkins, 2005; Zinoubi et al., 2017; Girardi et al., 2022) and swimming (Osiecki et al., 2014).^2^ Even in cycling, where the hyperbolic model and variations thereof dominate the literature, Coakley (2015) – see also Passfield et al. (2013) – suggested that

> “[the] power law is better than the [critical-power] model for predicting cycling endurance performance across a wider range of exercise intensities.” Coakley (2015, Chapter 5)

However, these studies do not appear to have had much impact on the above-mentioned debates surrounding the utility of the critical-power paradigm because they were purely empirical and based only on small sample sizes (between 8 and 15 individuals). In contrast, our evidence includes three large studies of 2571 runners, 3244 rowing ergometer seasons’ bests, and 5805 cyclists, as well as formal (i.e. mathematical) derivations.

## 2 Power–duration models

### 2.1 Setting

Power–duration models are functions which assert a relationship between an exercise duration *T* and a highest power *P*(*T*) which can be sustained over that duration. Throughout this work, we focus mostly on this *power–duration* relationship *P* = *P*(*T*). However, by simple algebra, this is always equivalent to, e.g., a particular *power–work* relationship *P* = *P*(*W*) which characterises the highest constant power which the athlete can continuously generate until some work *W* > 0 has been done; or to a particular *work–duration* relationship *W* = *W*(*T*) which gives the maximum amount of work that can be performed over duration *T*. All of these relationships can be inverted, e.g. *T* = *T*(*P*) is the time to exhaustion for some constant power output *P*.

Power meters are available on cycling and rowing ergometers and use of mobile power meters is now widespread in cycling, making it feasible to estimate and directly work with the power–duration relationship. However, for running, we will use “velocity” (*V*) as a proxy for “power” (*P*) and “distance” (*D*) as a proxy for “work” (*W*). This is commonly done in the literature and is justified as long as the conditions throughout the activity are relatively constant, e.g., flat course, no significant amount of wind, and negligible air resistance/friction.

We stress that *P*(*T*) (likewise *V*(*T*)) must be understood as the intensity that an athlete can *sustain* over a duration *T*. It does not model the initial acceleration (or the fatigue that occurs during the “acceleration phase”). Therefore, a cyclist’s observed maximal power output over one second will be below *P*(*T* = 1 s). Similarly, a runner’s observed average velocity in a 100-m sprint will be below *V*(*D* = 100 m). However, these effects become negligible over longer durations or distances.

### 2.2 Hyperbolic (a.k.a. critical-power) model

#### 2.2.1 Model

The *hyperbolic* model (Hill, 1925; Monod and Scherrer, 1965) (see also Jones et al., 2010, and references therein) assumes the following power–duration relationship which is also illustrated in Figure 1a:

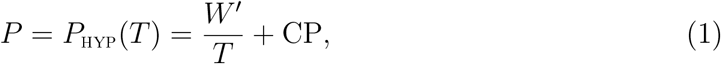

for parameters *W*′, CP > 0 and exercise durations *T* > 0. That is, (1) posits that athletes can continuously generate power *P*_HYP_(*T*) > CP for at most *T* seconds at which point they are exhausted. The parameters CP and *W*′ are interpreted as follows.

- **Critical power** CP. The additive constant CP > 0, called *critical power*, was originally interpreted as the power that could be sustained “for a very long time without fatigue” (Monod and Scherrer, 1965). However, this interpretation is no longer meaningful as the relationship in (1) is nowadays assumed to only be valid for exercise durations *T* that are not too long and not too short (we discuss this in Section 2.2.2 below).
- **Finite work capacity *W*′.** Multiplying both sides of (1) by *T* shows that athletes exercising at some constant power output *P* > CP will have generated a total amount of work equal to *W*′ + *T* · CP by the time they reach exhaustion. Thus, the parameter *W*′ is interpreted as representing a finite “tank” of exercise capacity “above CP” available to the athlete. As discussed below, this interpretation still holds even if the power output *P* > CP is not constant.

Simple algebra implies that the power–duration relationship from (1) (Figure 1a) can be equivalently written as a *power–work* relationship (Figure 1b) and *work–duration* relationship (Figure 1c). A full list of such equivalent relationships is given in Appendix A.1.1. Finally, as discussed in Section 2.1, we sometimes (e.g. in running) work with velocity (*V*) and distance (*D*) instead of power (*P*) and work (*W*). In this case, we write *D*′ instead of *W*′ and CV (*“critical velocity*”) instead of CP.

#### 2.2.2 Range of validity

The hyperbolic model, i.e. the relationship from (1), is nowadays assumed to only hold for exercise in the so-called “severe-intensity domain”, i.e. for exercise during which 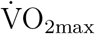 is attained (Jones et al., 2019). More formally, the hyperbolic model is assumed to be valid for exercise durations *T* such that *P*_L_ < *P*_HYP_(*T*) < *P*_U_, where *P*_L_ and *P*_U_ are the lower and upper boundaries of the severe intensity domain. Unfortunately, the requirement that the model should apply only to “severe” exercise is impractical. This is because this requirement typically leaves the range of validity of the hyperbolic model (i.e. the durations *T* or powers *P* for which the model is applicable) ill-defined for the following reasons:

- **Unclear boundaries.** The boundaries *P*_L_ and *P*_U_ of the “severe” domain differ across individuals and are typically unknown. As a result, it is not always clear which data points (i.e. which power–duration measurements or race results) one can include or exclude when fitting the model or even for which race distances the model is allowed to make performance predictions. For instance, it is not clear whether the model applies to a 5-km run or a 20-minute cycle ride.
- **Circular reasoning.** The requirement that the model should only be applied to “severe” exercise can also quickly become circular because the lower boundary of the severe domain, *P*_L_, is defined as being equal to the value of the parameter CP from the hyperbolic model, i.e. *P*_L_ ≔ CP. In other words, we should fit the hyperbolic model solely to exercise intensities above CP but we only know the value of CP *after* fitting the model to data. This circular reasoning is not merely a philosophical issue because the parameters *W*′ and CP of the hyperbolic model are highly sensitive to the range of durations (and powers) of the data points used. For instance, Gorostiaga et al. (2022b) point out that as we include data points over longer and longer durations, the estimate of CP decreases. The only way to avoid this circular reasoning is to estimate *P*_L_ = CP by other means but there is no simple alternative.^3^
- **Unbounded endurance.** Even if *P*_L_ and *P*_U_ were known exactly, assuming that the hyperbolic is valid for all durations *T* such that *P_L_* < *P*_HYP_(*T*) < *P*_U_ is still impractical. This is because the model would then formally imply that power outputs slightly above *P*_L_ = CP could be sustained “for a very long time without fatigue”. For instance, the model would predict that a cyclist with *P*_L_ = CP = 300 and *W*′ = 25 000 would be able to sustain 300.1 J/s for over 70 hours. However, this is no longer accepted in the literature (Vanhatalo et al., 2011; Poole et al., 2016).

For these reasons, we follow Jones et al. (2019) in assuming that the hyperbolic model is valid for exercise durations *T* between 2 and 15 minutes. We note that the 2–15 minute range is not viewed as being strict but rather as being a guide in the sense that exercise durations up to 20 or even 25 minutes can sometimes still be included. However, throughout this work, we will use 15 minutes, whenever possible, because:

1. Knowing that the model can *sometimes* still work over durations longer than 15 minutes does not lead to a practically useful rule for deciding to which efforts the model can or cannot be applied.
2. None of our findings would change qualitatively if we chose, say, 2–20 or 2–25 minutes. In fact, our results in the next section suggest that 2–15 minutes is actually the best-case scenario for the hyperbolic model: its errors (compared with the power-law model) quickly increase outside this range.

#### 2.2.3 Estimation

The parameters of the hyperbolic model are commonly estimated via linear regression by exploiting the fact that in (1), *P* is (affine) linear in 1/*T*. To keep the presentation simple we adopt this approach throughout this work (except for some of the panels in Figure 4). However, none of our results would change qualitatively if we used the more sophisticated non-linear weighted least-squares approaches (potentially based on the power–work relationship) advocated in Patoz et al. (2021). The estimation requires data from a sufficiently large number of maximal efforts (i.e. efforts to exhaustion) over different distances or durations with a time to exhaustion between 2 and 15 minutes (Karsten et al., 2015).

#### 2.2.4 Rate-of-exertion interpretation

Solving (1) for 1/*T* as in Gordon (2005) implies that, under the hyperbolic model, the athlete’s *rate of exertion* when generating some power *P* ≥ CP is given by

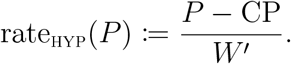

Thus, the accumulated exertion after exercise duration *t* is:

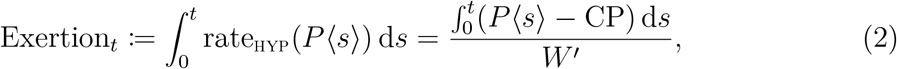

where *P*〈*s*〉 ≥ CP denotes the instantaneous power output at time *s*. That is, at the start of the activity, the athlete is assumed to be rested (Exertion_0_ = 0). The athlete is exhausted (and has to drop power output down to at most CP) at the first time *T* such that Exertion_*T*_ = 1.

Equation 2 shows that exhaustion occurs when the amount of work generated “above” CP, 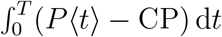, equals *W*′. Therefore, an often-used equivalent interpretation is that Balance_*t*_ ≔ 1 – Exertion_*t*_ ∈ [0, 1] is the remaining *balance* (measured as a proportion of *W*′) of the athlete, i.e. athletes start with a full “*W*′ tank” (Balance_0_ = 1) and they reach exhaustion at the first time *T* such that they have fully emptied this tank (Balance_*T*_ = 0).

### 2.3 Power-law (a.k.a. Riegel) model

#### 2.3.1 The model

The *power-law* model goes back to at least Kennelly (1906); Lietzke (1954) (see García-Manso et al. 2005; Vandewalle 2018 and references therein). It was popularised by Riegel (1977, 1981) – especially in the long-distance running community where it implies a widely used rule-of-thumb for finishing-time prediction (see, e.g., Runner’s World Magazine, 2013). It also forms the basis of the more sophisticated predictors from Blythe and Király (2016) and Vickers and Vertosick (2016). The latter is implemented by the websites Slate (Aschwanden, 2014) and FiveThirtyEight (Aschwanden, 2016).

The power-law model assumes the following power–duration relationship illustrated in Figure 2a. For parameters *S* > 0, 0 < *E* < 1 and any duration *T* > 0:

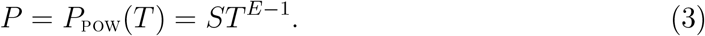

That is, (3) posits that athletes can continuously generate power *P*_POW_(*T*) > 0 for at most *T* seconds at which point they are exhausted.

As with the hyperbolic model, simple algebra allows us to convert (3) (Figure 2a) into an equivalent power–work relationship (Figure 2b) and work–duration relationship (Figure 2c). A full list of such equivalent relationships is given in Appendix A.2.1.

The rôle of the parameters *S* and *E* are visualised in Figure 3. Specifically:

- **Speed parameter *S***. The *speed* parameter *S* > 0 governs the vertical scaling of the power-duration curve. It is also the power that a cyclist can sustain for one second or the velocity that a runner can sustain for one second.
- **Endurance parameter *E*.** The *endurance* parameter 0 < *E* < 1 governs how quickly the power–duration curve in Figure 3 decays (smaller values of *E* correspond to a quicker decay). Its reciprocal, *F* = 1/*E* > 1, was termed *fatigue factor* by Riegel (1981). Throughout this work we will sometimes switch between *F* and *E* = 1/*F* to simplify the presentation; Riegel (1981) also gave suitable values for *F* in different sports and different demographic groups (derived from world-record performances *across* different athletes). We reproduce some of these in Table 1.

**Figure 3:**
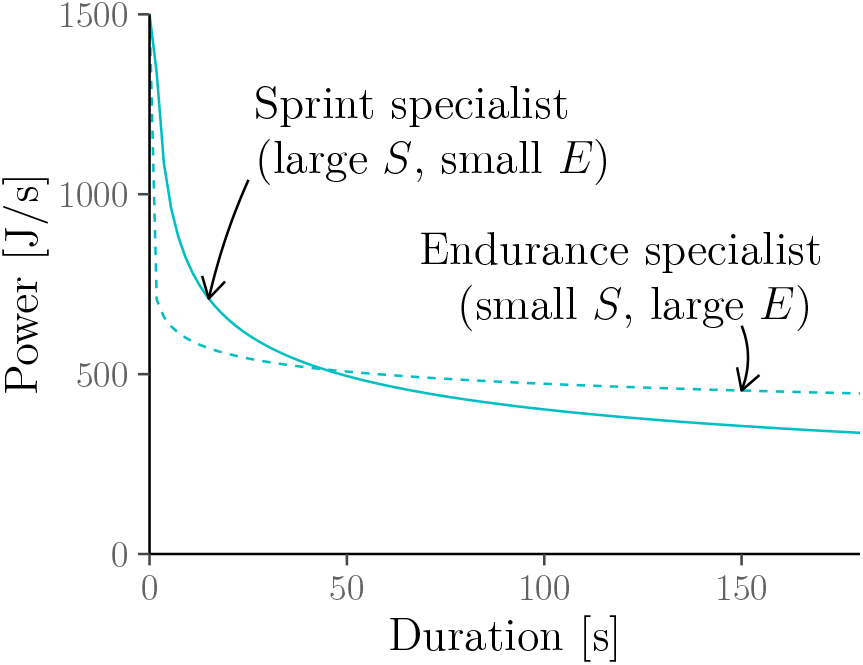
Rôle of the speed parameter *S* and the endurance parameter *E*.

**Table 1:** Fatigue factors derived from world records in Riegel (1981).

| Activity | Group | Fatigue factor $F$ | Endurance $E = 1/F$ |
| --- | --- | --- | --- |
| Running | Women | 1.08283 | 0.923506 |
|  | Men | 1.07732 | 0.928229 |
|  | Men (over 40) | 1.05352 | 0.949199 |
| Cycling | Men | 1.04834 | 0.953889 |

The power-law model has been independently discovered many times in the literature. For instance: Hinckson and Hopkins (2005); Laursen et al. (2007) applied it to running but called it the *log-log* model because it implies affine-linear relationships between the log-duration or log-work and log-power; Pinot and Grappe (2011) found it to perform well in cycling.

Finally, García-Manso et al. (2012); Blythe and Király (2016) have noted that there are systematic patterns in the residuals when fitting the power-law model to world-record performances in running. However, we stress that this considers the power–duration (or velocity–duration) relationship averaged *across* different athletes. In contrast, our work is concerned with modelling this relationship *within* individual athletes and we agree with Broxterman et al. (2022) that “[d]etermining the speed–duration relationship across athletes (…) while interesting, is fundamentally different from determining this relationship within an athlete”. Indeed, Blythe and Király (2016) found that the power-law model performs better for the latter than for the former based on 1 417 432 performances of 164 746 runners (a subset of which we analyse in the next section):

> “(…) although world-record performances are known to obey a power law (…), there is no reason to suppose a-priori that the performance of individuals is governed by a power law. Striking is that the power-law derived is considerably more accurate when considered in log-distance–log-speed coordinates than the power-law which applies to world-record data.” (Blythe and Király, 2016)

#### 2.3.2 Range of validity

As we will show in the next section, the power-law model describes exercises capacity well across a wide range of durations. In particular, unlike the hyperbolic model, the power-law model is not restricted to predicting only those power outputs that fall into a particular range (e.g. into a particular exercise-intensity domain). Note that this makes the power-law model much easier to use and more widely applicable than the hyperbolic model. Of course, there are some obviously “silly” edge cases that should be avoided such as applying the model to durations *T* longer than an individual’s lifespan. We also reiterate our comment from Section 2.1 that observed performances over short efforts (e.g. 100m sprints) will be below what the model predicts because the model describes the power or velocity that can be *sustained* over a particular duration or distance and thus does not account for the initial acceleration (whose impact in short races is substantial).

#### 2.3.3 Estimation

The power-law model can again be easily estimated via linear regression by exploiting the fact that by (3), log(*P*) is (affine) linear in log(*T*). In this case, *exp(intercept*) is an estimate of *S* and (*slope* + 1) is an estimate of *E*. To keep the presentation simple, we adopt this approach throughout this work. However, none of our results would change qualitatively if we used linear regression based on the fact that log(*P*) is (affine) linear in the log-work log(*W*) or if we used more sophisticated non-linear least-squares approaches such as those advocated for the hyperbolic model in Patoz et al. (2021).

If not enough data points are available to accurately estimate both parameters, e.g. when predicting a marathon finish time from a single recent half-marathon result, it can be preferable to fix *E* (equivalently: *F* = 1/*E*) to a suitable default value, e.g. to one of the values provided in Table 1. For instance, consider the popular calculator from Runner’s World Magazine (2013) which predicts the finish time *T* in a race over distance *D* from the finish time *T*_0_ in a previous race over some other distance *D*_0_ as

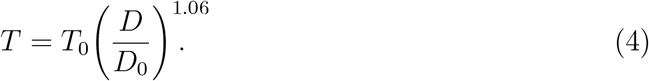

This calculator is obtained from (3) by fixing *F* = 1.06 (see Appendix A.2.2 for details).

Of course, the endurance parameter *E* (equivalently: the fatigue factor *F*) is likely to be different for different athletes as shown in Blythe and Király (2016); Zinoubi et al. (2017). Indeed, if it was not, we could expect some athletes to simultaneously hold world records over both short and long race distances. Hence, setting *E* to a default value incurs a bias. However, this bias is outweighed by the variance reductions brought about by avoiding the need for estimating *E* from limited amounts of data. Thus, fixing the fatigue factor to a sensible default value can be justified as a *bias–variance trade-off*.

#### 2.3.4 Rate-of-exertion interpretation

We end this section by showing that we can derive a similar “rate-of-exertion” interpretation for the power-law model as for the hyperbolic model. Solving (3) for 1/*T* implies that the athlete’s *rate of exertion* when generating some power *P* ≥ 0 implied by the power-law model is given by

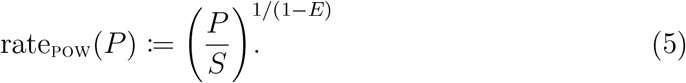

Thus, the accumulated exertion after exercise duration *t* is:

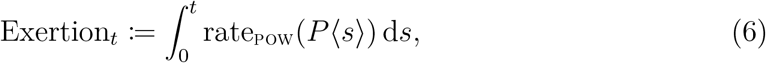

where *P*〈*s*〉 ≥ 0 denotes the instantaneous power output at time *s*. That is, at the start of the activity, the athlete is assumed to be rested (Exertion_0_ = 0). The athlete is exhausted (and has to drop power output down to 0) at the first time *T* such that Exertion_*T*_ = 1.

Again, we can equivalently interpret Balance_*t*_ ≔ 1 – Exertion_*t*_ ∈ [0, 1] as a remaining “balance” (as a proportion of 1/*S*^1/(1–*E*)^) of the athlete. Importantly, (5) shows that this balance decreases faster than linearly in the power output, *P*. This is in contrast to the hyperbolic model whose balance decreases only linearly in *P* (more accurately: linearly in *P* – CP). For this reason, no finite work capacity (i.e. “*W*′-tank”) interpretation is possible for the power-law model.

To our knowledge, the rate-of-exertion interpretation of the power-law model is novel. In Section 4, we employ this construction to prove a mathematical statement which suggests that the power-law model has more realistic pacing implications than the hyperbolic model. In Section 5, we employ this construction to show that, consistent with empirical evidence, the power–duration curve implied by the power-law model moves downwards as the athlete becomes more fatigued.

## 3 Contribution I: Fit over different durations

### 3.1 Overview

It is well known that the hyperbolic model adequately describes exercise capacity over durations that are not too short and not too long, e.g. around 2–15 minutes, but exhibits unrealistic behaviour outside this range (see, e.g., Vandewalle et al., 1997; Jones et al., 2019; Pallarés et al., 2020).

In this section, we empirically show that the good fit of the hyperbolic model for exercise durations in the 2–15 minute range does not mean that the power–duration relationship is hyperbolic. Rather, the power–duration relationship can be much better described by a power law and the hyperbolic model just happens to approximate the power-law model reasonably well in the 2–15 minute range.

We first demonstrate this result in the context of the case studies of elite runners from Jones et al. (2019); Jones and Vanhatalo (2017). We then show similar results in large-data studies of recreational runners, rowers and cyclists. Finally, we explain why the problems of the hyperbolic model become even more apparent when combining it with a different functional form over long durations, e.g., as done in the so-called *omni power–duration* models from Puchowicz et al. (2020).

### 3.2 Case study from Jones et al. (2019)

Here, we reproduce the case study of Eliud Kipchoge from Jones et al. (2019). We fit the hyperbolic model to Eliud Kipchoge’s personal records for races in the 2–15 minute range (*D*′ ≈ 176.29 m, CV ≈ 6.23m/s) and compare the results with the power-law model fitted to all of Eliud Kipchoge’s personal records in different events (including indoor events) up to marathon distance (*S* ≈ 9.57m/s, *E* ≈ 0.94). ^4^ The data was obtained from World Athletics (2021). For completeness, we provide Kipchoge’s personal records, e.g., 00:13:11 (5km); 00:59:25 (half marathon) and 02:01:39 (marathon), in Appendix B.2.

The results are shown in Figure 4 which replicates all four panels from Jones et al. (2019, Figure 5). It illustrates that, for Eliud Kipchoge’s data, the hyperbolic model predicts unrealistically high velocities for durations shorter than 2 minutes or longer than 15 minutes. In contrast, the power-law model still predicts realistic velocities over a wide range of durations.

- **Short durations.** If we were to apply the hyperbolic model to shorter exercise durations, it would imply that Eliud Kipchoge can
  – sustain more than 650km/h over 1 second (85km/h over 10 seconds);^5^
  – instantaneously “teleport” over *D*′ ≈ 176 metres (see the non-zero intercept in Figure 1c; Proposition 3 in Appendix B.1 gives a formal proof);
  In contrast, the power-law model implies that Eliud Kipchoge can
  – only sustain *less than* 35km/h over 1 second (30km/h over 10 seconds) – well below the top speeds of elite sprinters;
  – *not* cover any non-zero distance instantaneously (see the zero intercept Figure 2c; Proposition 4 in Appendix B.1 gives a formal proof).
- **Long durations.** If we were to apply the hyperbolic model to longer exercise durations, it would imply that Eliud Kipchoge can run ultra-marathons at essentially the same average velocity as half-marathons. In contrast, the power-law model implies that the velocity which Eliud Kipchoge can sustain over some duration/distance goes to zero as the duration/distance increases.

**Figure 4:**
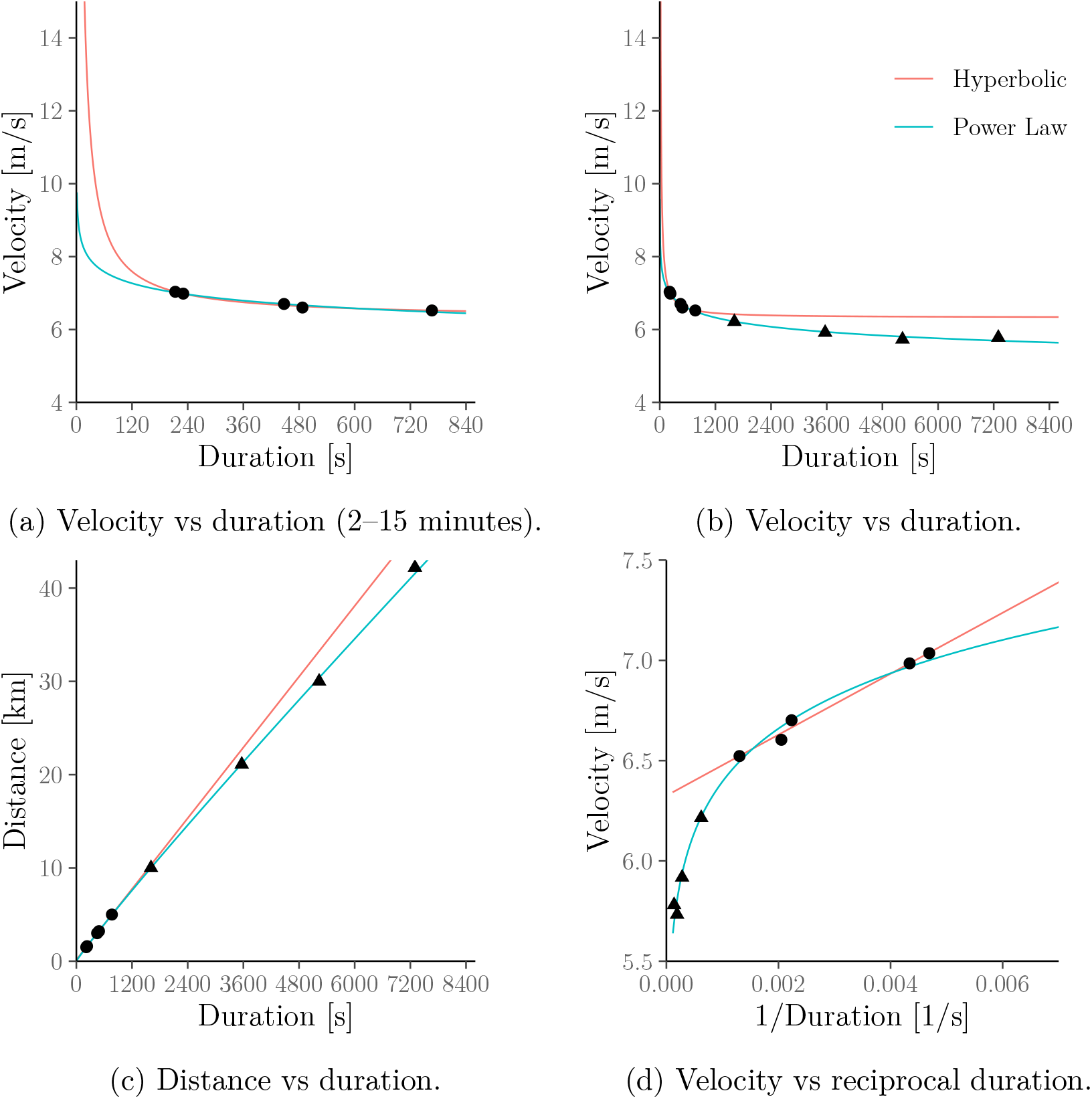
An extended and modified version of Jones et al. (2019, Figure 5, Panels A–D) showing the hyperbolic (a.k.a. critical-power) model fitted to Eliud Kipchoge’s personal records. As in Jones et al. (2019), the hyperbolic model is fitted only to the personal records in the 2–15 minute range (●); the power-law (a.k.a. Riegel) model is fitted to all personal records up to marathon distance (● & ▲). Note that the power-law model behaves similarly to the hyperbolic model inside the 2–15 minute range but exhibits more realistic behaviour outside this range.

**Figure 5:**
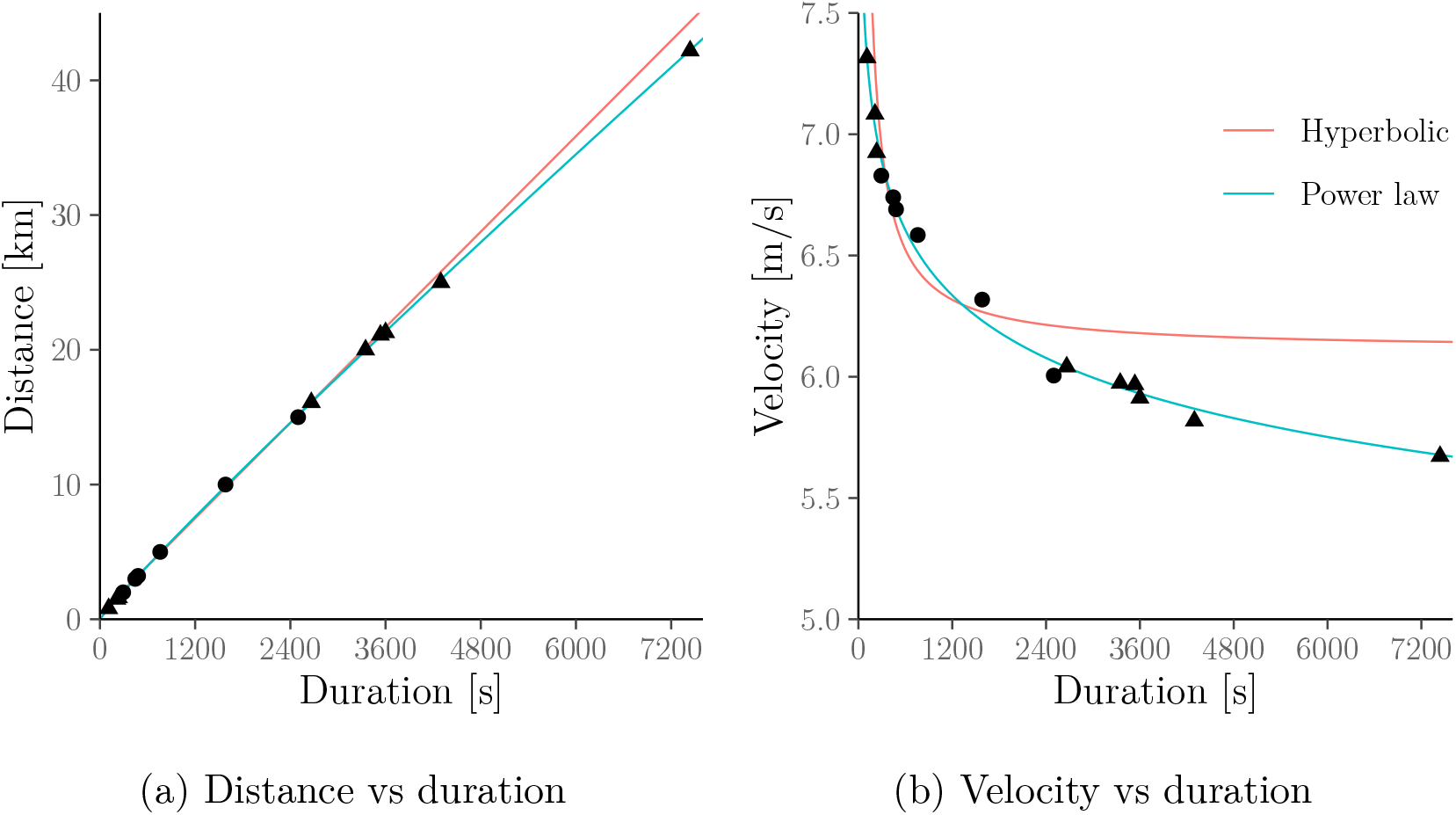
Haile Gebrselassie’s personal records in different events. Figure 5a extends Jones and Vanhatalo (2017, Figure 2) to include races shorter than 1500m and longer than 15 000m. As in Jones and Vanhatalo (2017) the hyperbolic (a.k.a. critical-power) model is only fitted to the personal records in the 1500–15 000m range (●); the power-law (a.k.a. Riegel) model is fitted to all personal records up to marathon distance (● & ▲). The figure shows that the power-law model exhibits realistic behaviour over all durations up to marathon distance.

Interestingly, in Figure 4, the hyperbolic and power-law models give very similar behaviour for exercise durations in the 2–15 minute range. Later in this section, we show that this holds for wider populations.

### 3.3 Case study from Jones and Vanhatalo (2017)

We now reproduce the case study from Jones and Vanhatalo (2017), i.e. we fit the hyperbolic model to the personal records over different events (including indoor events) up to marathon distance of Haile Gebrselassie and 11 other elite runners. The data were again obtained from World Athletics (2021).

We apply the hyperbolic model to all races with distances between 1500m and 15 000m which corresponds to exercise durations of around 3.6 to 41.6 minutes for Haile Gebrselassie. Clearly, this does not match the 2–15 minute exercise duration range for which the hyperbolic model is thought to be valid but is done to match the analysis carried out in Jones and Vanhatalo (2017). We compare the results with the power-law model (fitted to all personal records up to marathon distance).^6^ For completeness, we provide Gebrselassie’s personal records, e.g., 00:12:39 (5000m); 00:58:55 (half marathon) and 02:03:59 (marathon), in Appendix B.3.

The first panel in Figure 5 extends Jones and Vanhatalo (2017, Figure 2) and shows that the power-law model fits well for Haile Gebrselassie’s personal records over different durations. For clarity, we have added the second panel which shows the same results but using the velocity–duration relationship.

Figure 6 compares the fit of both models for the group of 12 male elite athletes (which include Eliud Kipchoge and Haile Gebrselassie) considered in Jones and Vanhatalo (2017, Table 1). A more formal numerical comparison of the errors of both models is given in Table 4 in Appendix B.3.

**Figure 6:**
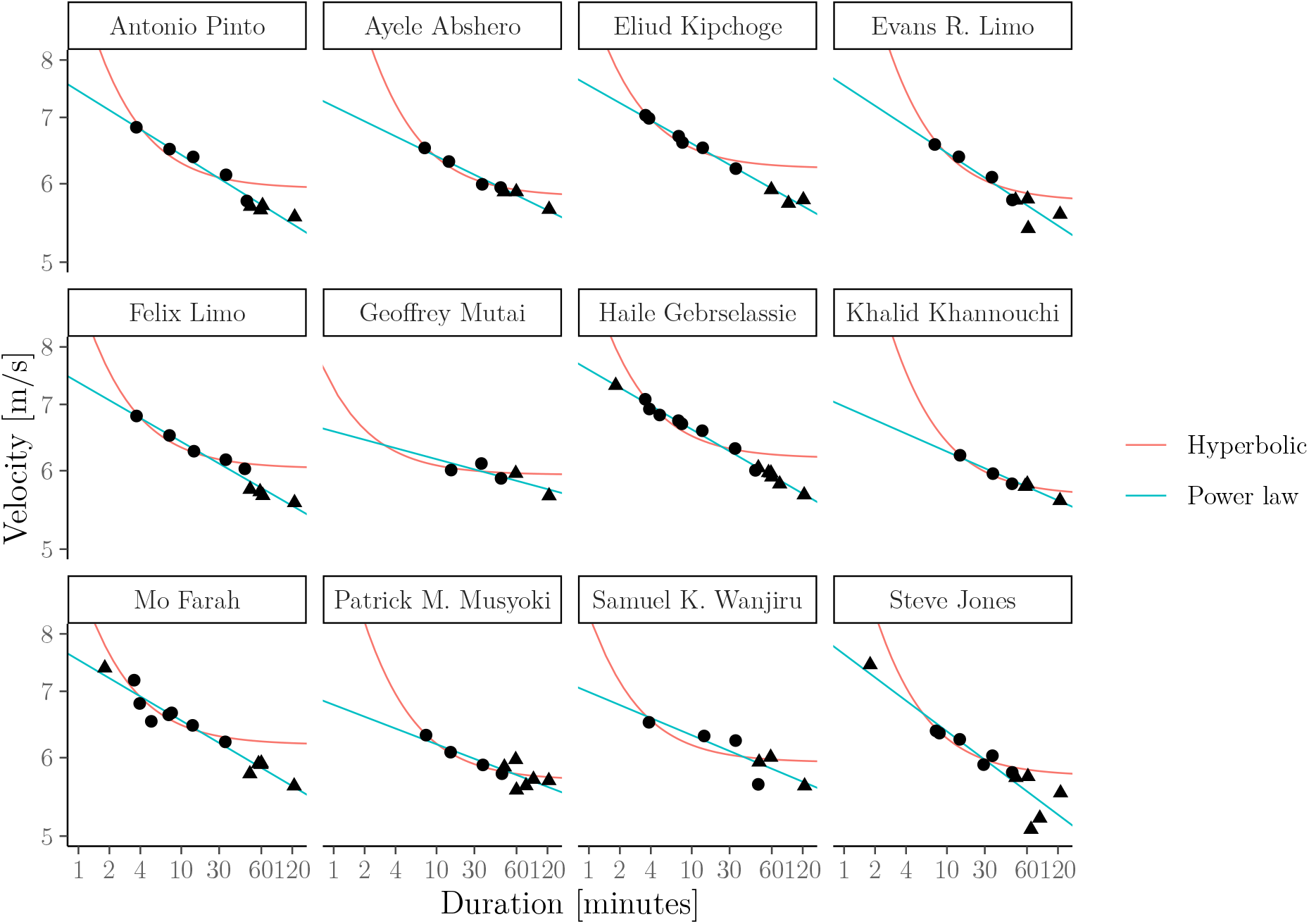
Personal records of the elite runners considered in Jones and Vanhatalo (2017, Table 1). As in Jones and Vanhatalo (2017) the hyperbolic (a.k.a. critical-power) model is only fitted to data in the 1500–15 000m range (●); the powerlaw model (a.k.a. Riegel) is fitted to all personal records up to marathon distance (● & ▲). Note that both axes are on the log-scale for clarity. The figure again shows that the power-law model exhibits realistic behaviour over all durations up to marathon distance (see Table 4 in Appendix B.3 for a more formal comparison of the fit of each model).

### 3.4 Large-data study in running

In the remainder of this section, we show that the above findings generalise to wider populations. First, we fit both models to race results from (mostly recreational) runners collected by Blythe and Király (2016) from the database `powerof10.info`. After removing athletes with too few records in the 2–15 minute range (see Appendix B.5 for details) we are left with race results for 2571 runners. For each of these, we fit both the hyperbolic and power-law models and compare the average error across all athletes, where errors defined as the average relative difference between observed and predicted velocities as in Puchowicz et al. (2020) (see Appendix B.4). Figure 7 shows this error averaged across all athletes for each model. This error is similar for both models for exercise durations between 2 and 15 minutes but worse for the hyperbolic model if we include efforts shorter than 2 minutes or longer than 15 minutes.

**Figure 7:**
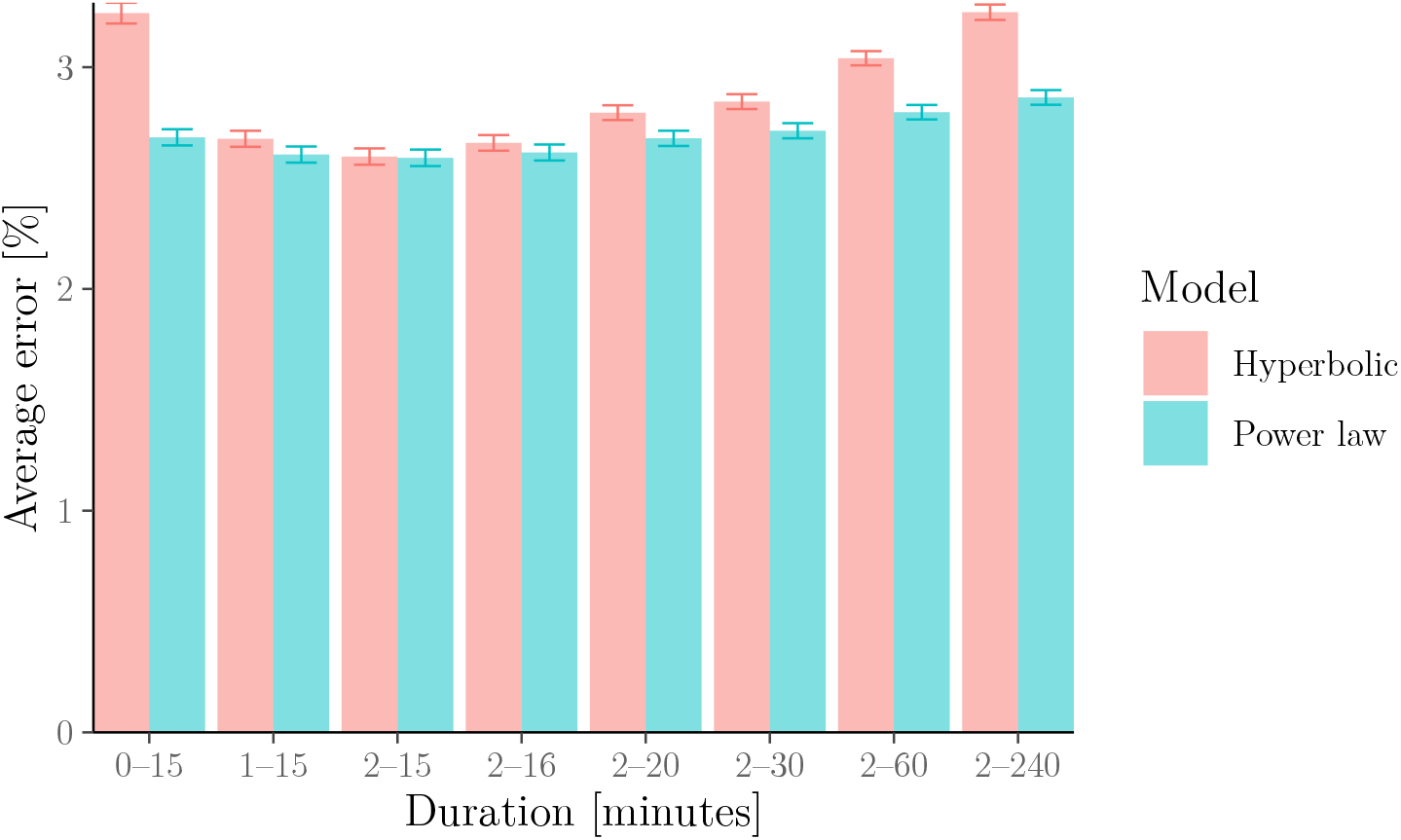
Relative error of the hyperbolic (a.k.a. critical-power) model vs the power-law (a.k.a. Riegel) model for 2571 runners. Errors are calculated as observed vs predicted relative velocities. The error bars represent standard errors. The figure illustrates that both models provide a similar fit for exercise durations in the 2–15 minute range but that the power-law model provides a better fit outside this range.

### 3.5 Large-data study in rowing

Next, we fit both models to rowing-ergometer data available from www.nonathlon.com. Specifically, the data contain self-reported seasons’ bests (for seasons from 2002 to 2022) in the form of finish times over fixed distances 500m, 1km, 5km, 6km, 10km, 21.0975km (half marathon) and 42.195km (marathon) as well as distances covered over fixed durations 30 min and 60 min. In total, we are left with 3244 athlete seasons after removing all athlete seasons with less than three measurements between 2 and 20 minutes (see Appendix B.6 for details). We stress that we selected 20 minutes instead of 15 minutes as the minimum upper bound on the duration ranges because (a) only 506 athletes in the data set have at least three efforts in the 2–15 minute range; (b) none of these have efforts shorter than 2 minutes. (c) all of these are “slow” athletes whose time over 500m is above 2 minutes (i.e. focusing only on these athletes might incur a selection bias).

We again fit both models to the data. The results are shown in Figure 8 which again illustrates that the power-law model describes the data better than the hyperbolic model over a wide range of durations.

**Figure 8:**
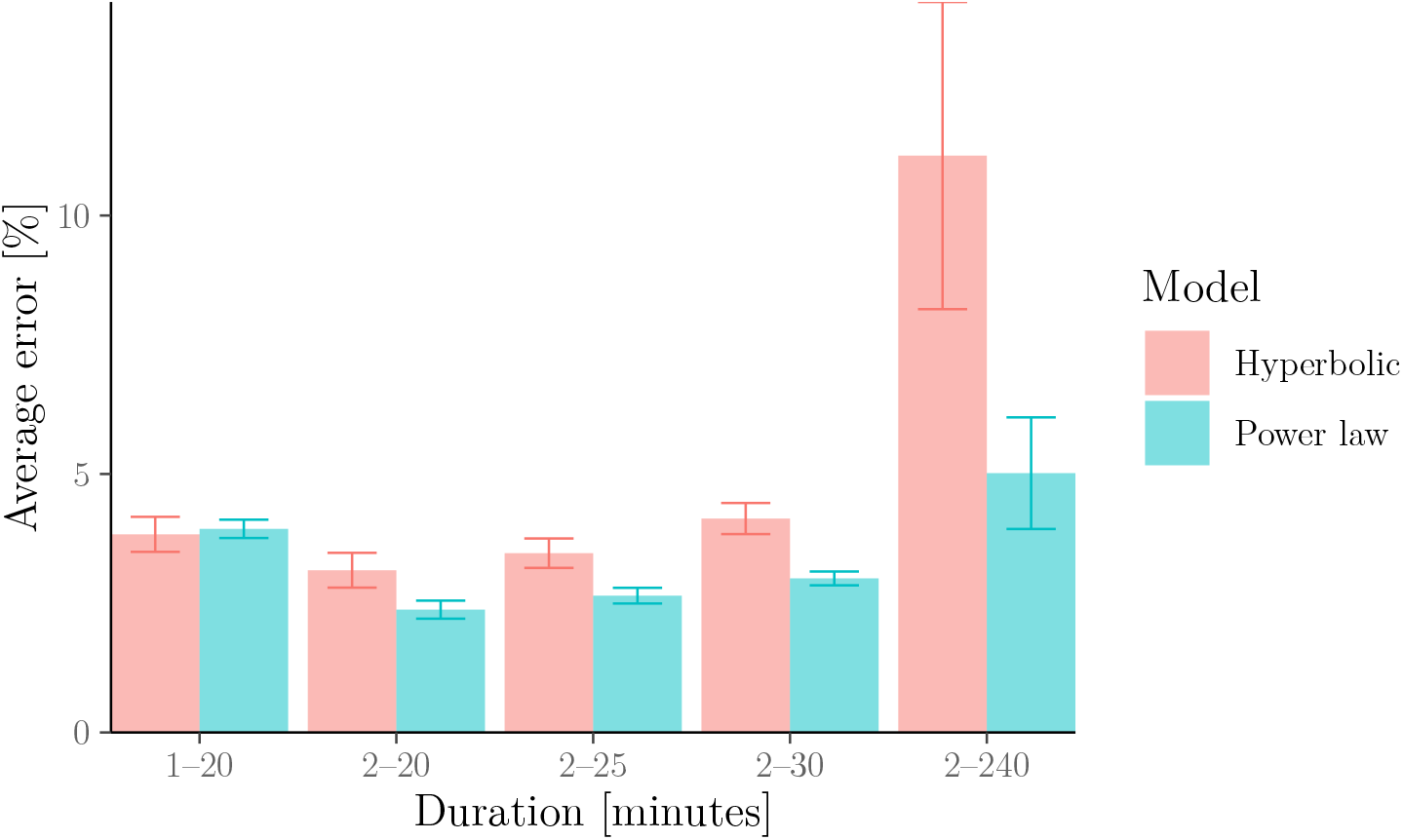
Relative error of the hyperbolic (a.k.a. critical-power) model vs the power-law (a.k.a. Riegel) model for 3244 athlete seasons in rowing. The errors and error bars are calculated as in Figure 7. The figure again illustrates that the powerlaw model describes the data more adequately than the hyperbolic model over a wide range of durations longer than 2 minutes (note that results for 1–20 minutes are inconclusive due to the overlapping error bars).

### 3.6 Large-data study in cycling

We now show (in Figure 9) the same analysis as in Section 3.4 but applied to an open data set of 5805 cyclists from Golden Cheetah (www.goldencheetah.org). For each athlete, we extracted the best average power output for a range of different durations from their training and racing history. It is well known that it is difficult to fit these models on racing and training data as it is unclear whether the data corresponds to truly maximal efforts (Puchowicz et al., 2018; Leo et al., 2021, 2022b). To alleviate this, we removed some efforts that could not have been maximal as well as outliers which are likely to be due to power-meter malfunctions (see Appendix B.7 for more details).

**Figure 9:**
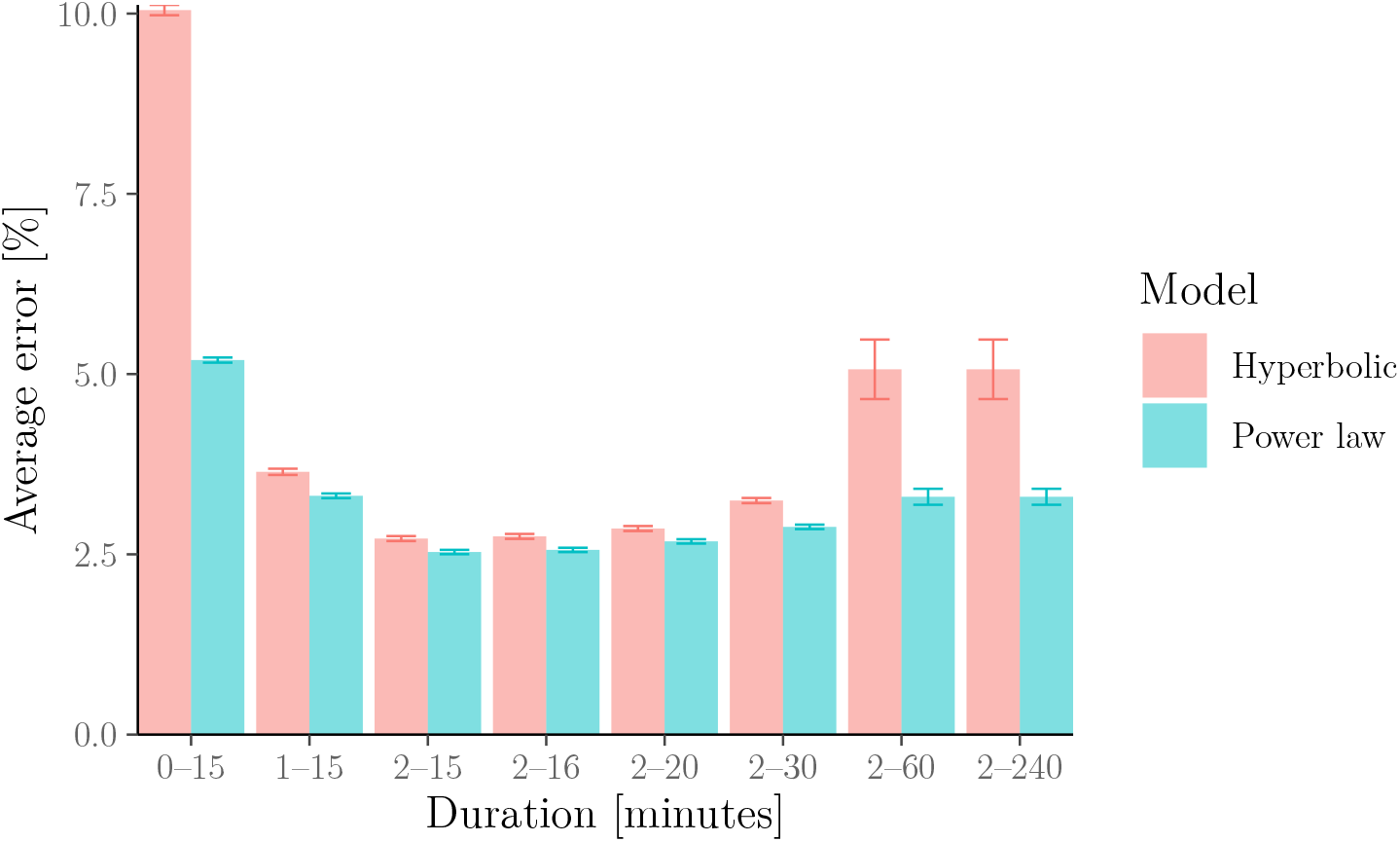
Relative error of the hyperbolic (a.k.a. critical-power) model vs the power-law (a.k.a. Riegel) model for 5805 cyclists. Errors are calculated as observed vs predicted relative powers. The error bars represent standard errors. The figure shows that errors are similar for exercise durations in the 2–15 minute range but that the power-law model has smaller errors than the hyperbolic model outside this range.

### 3.7 Piecewise-defined models

To permit more realistic predictions outside the 2–15 minute range without abandoning the critical-power paradigm, it has been proposed to combine (variants of) the hyperbolic model for durations shorter than some threshold *T*_*_ > 0 with different functional forms for durations longer than *T*_*_ (Péronnet and Thibault, 1989; Puchowicz et al., 2020; Luttikholt and Jones, 2022) – see Figure 10, and Appendix B.8 for a more formal treatment. However, such “piecewise-defined” models introduce other drawbacks:

1. They require an increased number of unknown parameters so that a large number of data points (e.g. from arduous critical-power tests or previous races) are needed to avoid overfitting – and a number of these data points must be from efforts longer than *T*_*_. Additionally, the threshold *T*_*_ is difficult to choose but its choice affects the other model parameters, e.g. CP and *W*′ (Puchowicz et al., 2020).
2. They posit a power–duration relationship that is actually no longer hyperbolic for exercise intensities above CP (or above similar parameter), i.e. they are not compatible with the assumption that athletes have a fixed amount of work, *W*′, that they can generate above CP.
3. They imply a power–duration relationship with unrealistic properties:
  a. For sufficiently long durations, the power–duration curve from Péronnet and Thibault (1989); Puchowicz et al. (2020) can become negative.
  b. The models from Péronnet and Thibault (1989); Puchowicz et al. (2020); Luttikholt and Jones (2022) have a “kink” in the power curve as shown in Figure 10.

**Figure 10:**
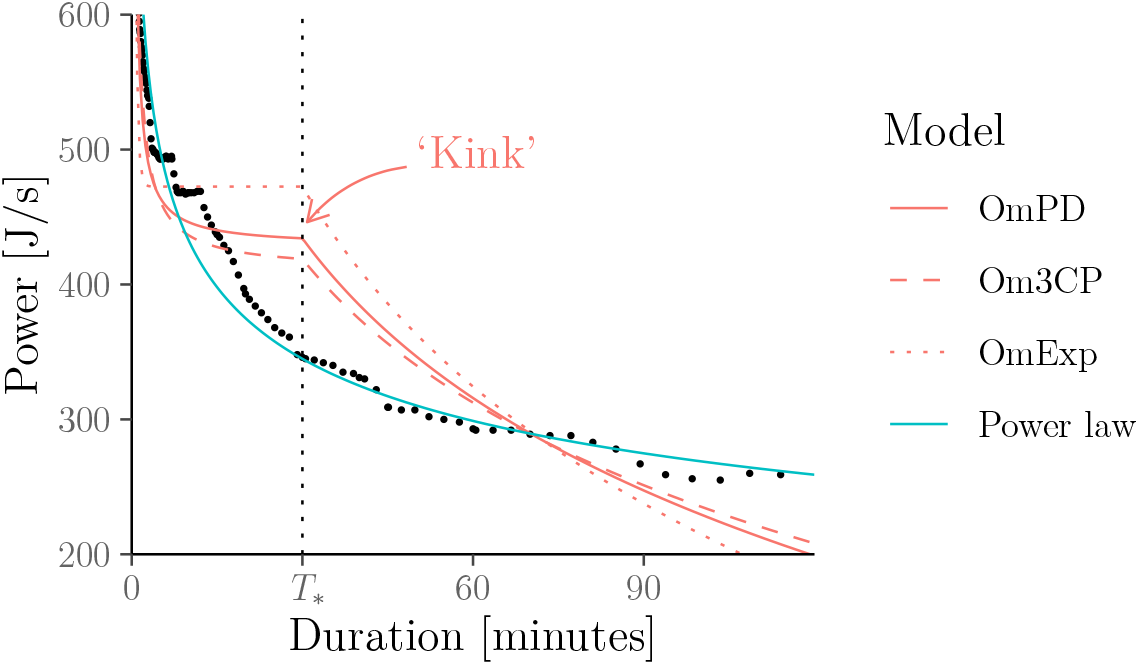
Power–duration relationship of the so-called omni power–duration models (OmPD, Om3CP, and OmExp; with the same threshold *T*_*_ = 30 min) from Puchowicz et al. (2020) for the data from Leo et al. (2022a, Figure 4) (●).

The last point, 3b, connects to our findings from earlier in this section: The hyperbolic model fits power–duration (or similar) data reasonably well for exercise durations *T* which are not too short (e.g. at least 2 minutes) and not too long (e.g. longer than *T*_*_ = 15 minutes). However, as we saw, this is only because the hyperbolic function approximates a power law reasonably well in this range. Indeed, the fact that the hyperbolic assumption is problematic becomes immediately apparent in Figure 10. Here, the need for reconciling the posited hyperbolic shape over durations *T* ≤ *T*_*_ with the fact that the power–duration curve should approach zero as *T* increases (as no exercise intensity can be maintained “for a long time without fatigue”) causes a “kink”.

While this kink could be easily smoothed out, this would still leave a power–duration curve with turning/inflection points.^7^ Indeed, unless *T*_*_ is trivially small, *any* model which combines a hyperbolic curve (below *T*_*_) with another curve that decays to zero for long durations (above *T*_*_) must exhibit such a kink, turning/inflection point, or discontinuity; and to our knowledge, there is no empirical evidence for the existence of such artefacts.^8^ This again calls the “hyperbolic” assumption into question.

## 4 Contribution II: Implications for pacing

### 4.1 Overview

In this section, we show that the power-law (a.k.a. Riegel) model has realistic pacing implications in the sense that it implies that overpacing (e.g. running or riding off too quickly) is detrimental to the overall finish time in a race. This is in contrast to the hyperbolic (a.k.a. critical-power) model which implies that the safest optimal pacing strategy is to sprint off as fast as possible and then “hold on”.

### 4.2 Setting

Throughout this section, we assume that we want to generate some power *W* over the shortest possible duration. Consider the following constant (“even”) pacing and variable (“uneven”) pacing strategies:^9^

**Pace_con_** Generate power *P* > 0 continuously over duration *T* > 0 such that exhaustion occurs at time *T* at which point work *W* has been accumulated, i.e. *T*· *P* = *W*.
**Pace_var_** Generate power *P*_1_ > 0 until some work 0 < *W*_1_ < *W* has been accumulated, i.e. maintain *P*_1_ over the duration *T*_1_ ≔ *W*_1_/*P*_1_. Subsequently, maintain the highest possible power *P*_2_ > 0 which allows further work *W*_2_ ≔ *W* – *W*_1_ to be accumulated, i.e. maintain *P*_2_ over the duration *T*_2_ ≔ *W*_2_/*P*_2_, where *P*_2_ is chosen such that exhaustion occurs when work *W* = *W*_1_ + *W*_2_ has been accumulated, i.e., at time *T*_1_ + *T*_2_.

Recall that under the assumptions made in Section 2.1, we can replace “work” by “distance” and “power” by “velocity”. For instance, when running a 5-km race, the Strategy **Pacecon** implies running continuously at the highest possible *constant* velocity that can be sustained over *D* = 5000m; Strategy **Pace_var_** implies running parts of the distance at different velocities, e.g. running the first *D*_1_ = 1000m at some velocity *V*_1_ and then running the remaining *D*_2_ = *D* – *D*_1_ = 4000m at the velocity *V*_2_ which is chosen such that exhaustion occurs when crossing the finishing line.

### 4.3 Impossibility of overpacing under the hyperbolic model

Under the hyperbolic model, any pace selection which ensures that (a) *W*′ is completely depleted by the end of the activity (see Section 2.2.4); (b) the power output never drops below CP, is optimal. In principle, athletes could just “burn through” their *W*′ tank (see Section 2.2.4) in the first second of the activity and then complete the rest of the activity at CP. In other words, overpacing is impossible in the hyperbolic model; athletes only ever need to worry about “underpacing” i.e. accidentally dropping the power output below CP. Thus, since CP is unlikely to be known exactly to the athlete, the hyperbolic model implies that the safest optimal pacing strategy is starting as fast as possible (to avoid the risk of dropping the power output below CP) and then simply “holding on”.

This result was already proved in Fukuba and Whipp (1999). For completeness, Proposition 1 (proved in Appendix C.1) repeats their result here in the setting described above. It shows that in the hyperbolic model,^10^ Strategies **Pace_con_** and **Pace_var_** both lead to the same optimal finishing time as long as *P*_1_ ≥ CP, i.e. as long as the athlete does not start out too slowly under **Pace_var_**. The solid red line in Figure 11 visualises Proposition 1 in the context of a hypothetical 5-km race (i.e. with power replaced by velocity and work replaced by distance – see Section 2.1).

**Figure 11:**
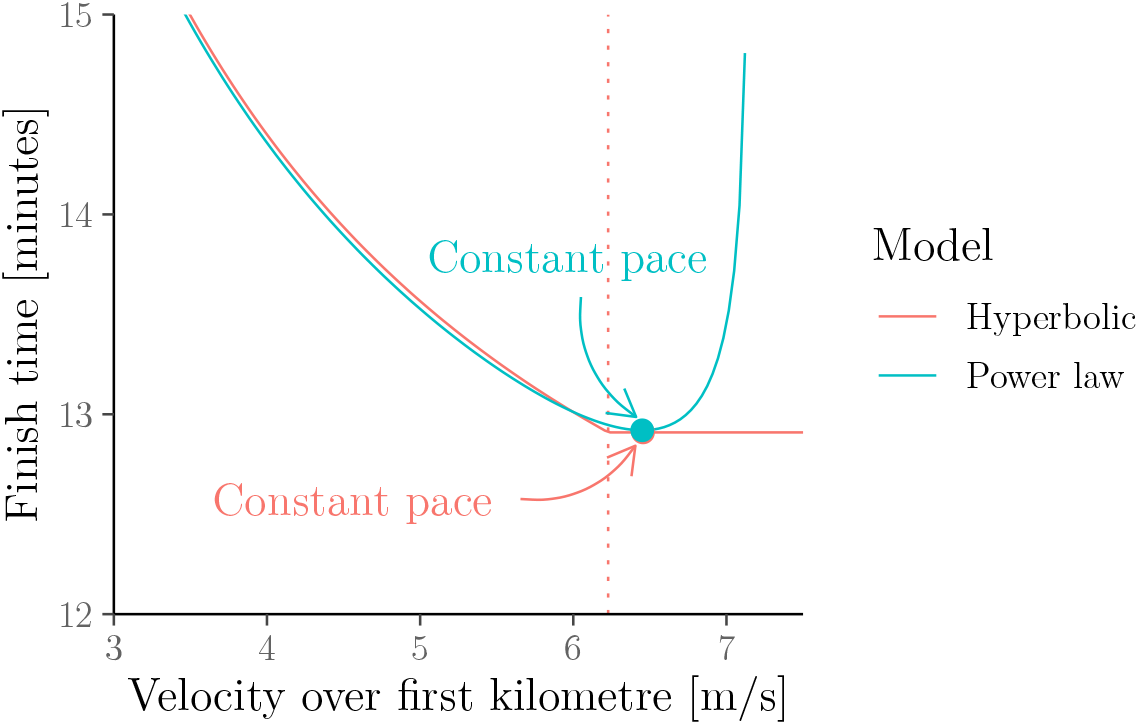
Illustration of Propositions 1 and 2 on a hypothetical 5-km race by Eliud Kipchoge. The lines show Kipchoge’s best possible finish times according to the hyperbolic (a.k.a. critical-power) and power-law (a.k.a. Riegel) model if he maintains the velocity given on the first axis over the first kilometre. The model parameters are as estimated in Section 3.2; the dotted line represents critical velocity. The figure illustrates that the power-law model considers overpacing to be dangerous whilst the hyperbolic model does not.

#### Proposition 1

(Fukuba and Whipp 1999). *For some work W* > *W*′, let *T, P* as well as (*P_i_, T_i_, W_i_*) for *i* ∈ {1, 2} be as *in Strategies* **Pace_con_** *and* **Pace_var_**, *with P*_1_ ≠ *P, then*:

1. *if P*_1_ ≥ CP, *then T*_1_ + *T*_2_ = *T*;
2. *if P*_1_ < CP, *then T*_1_ + *T*_2_ > *T*.

### 4.4 Optimality of even pacing under the power-law model

We now present our main result in this section: Proposition 2 (proved in Appendix C.2) shows that in the power-law model, the “even pacing” strategy, **Pace_con_**, yields better finishing time than the “uneven pacing” strategy, **Pace_var_**. That is, pacing is crucial in the power-law model: any deviation from the optimal constant pace leads to an increase in the finishing time (and overpacing is worse than “underpacing”). The blue line in Figure 11 visualises the result of Proposition 2. To our knowledge, Proposition 2 and its proof (which which uses our new “rate-of-exertion” interpretation of the power-law model presented in Section 2.3.4) are novel.

#### Proposition 2.

*For some work W* > 0, *let T, P as well as* (*P_i_, T_i_, W_i_) for i* ∈ {1, 2} *be as in Strategies* **Pace_con_** *and* **Pace_var_**. *Then the finish time under* **Pace_var_**, *T*_1_ + *T*_2_, *grows as the difference between the initial and the optimal even power output*, |*P*_1_ – *P*|, *increases. In particular, T*_1_ + *T*_2_ > *T whenever P*_1_ ≠ *P*.

### 4.5 Pacing results in context

We end this section by putting the pacing implications of the hyperbolic and power-law models in the context of existing research.

To ensure that the goal of “winning a race” can be used as a proxy for the goal of “minimising the amount of time needed to generate a fixed amount of work” which we analysed above, we restrict our attention to, essentially, time trials that have standard conditions, e.g. constant course topography and insignificant wind speeds, constant temperature/humidity, and the absence of tactics/drafting or other psychological factors that might interact with pacing. We also assume that the race is long enough such that the impact of initial acceleration or the waste of kinetic energy at the end of the race is negligible (de Koning et al., 2011).

Under these assumptions, it is thought that the optimal performance in a race is achieved through an even-pacing strategy (Abbiss and Laursen, 2008). For instance:

- **Running.** In long-distance running, even pacing is optimal under models based around physics (Keller, 1974). Additionally, the previously mentioned online running finish-time predictors (e.g., Runner’s World Magazine, 2013; Aschwanden, 2014) are motivated by the idea an even pace is critical to achieving the best possible performance for a given fitness level. Empirically, faster finishing times are also positively correlated with more even pacing strategies (e.g., Hoffman, 2014; Keogh et al., 2020). Finally, even pacing is also recommended by most coached (e.g., Galloway, 2007; Pfitzinger and Douglas, 2009) and observed in elite athlete performances. For instance, the 5-km-split times in Eliud Kipchoge’s sub-2-hour marathon in 2019 were 00:14:12 (±2 seconds).
- **Cycling.** In cycling, even pacing strategies are considered to be favourable under the assumptions made above (Foster et al., 1993; de Koning et al., 1999; Atkinson and Brunskill, 2000; Atkinson et al., 2003; Ham and Knez, 2009); see Coakley and Passfield (2018) for a review and further references. Under the controlled conditions of a 60-minute world record attempt, the cyclist in Padilla et al. (2000, Figure 2) chose a relatively even pace.

We stress again that the assumptions made in this section, and by extension, the optimality of even pacing, may not hold in very short races where the impact of initial acceleration and wasted kinetic energy at the end as well as of other, e.g. physiological or psychological factors, are non-negligible. For instance:

- In a physics-based model form de Koning et al. (1999), fast starts are optimal in 1000m track cycling events. However, in longer, e.g. 4000m events, it is optimal to transition to even pacing after a fast start over the first few seconds.
- Bishop et al. (2002) observed that fast starts over the first few seconds (again followed by a transition to constant pace) correlated with improved performance in two-minute kayak ergometer tests.
- Fast starts were observed to correlate with improved performance in three-minute efforts in a study of seven cyclists from Bailey et al. (2011). However, this effect was not observed in six-minute efforts.^11^

In summary, the literature appears to be consistent with the power-law model’s implication that athletes should implement an even pacing strategy (except during the initial acceleration phase which can make up a significant proportion of the effort during short races, e.g. sprints) but not consistent with the hyperbolic model’s implication that athletes never need to worry about overpacing (all under the assumptions made above).

## 5 Contribution III: Modelling fatigue

### 5.1 Overview

It has been found that the power–duration curve moves downwards as prolonged exercise causes the athlete to fatigue (Leo et al., 2021). This behaviour is not captured by the hyperbolic (a.k.a. critical-power) model as shown in Figure 12. To circumvent this problem, it has been suggested that CP decreases with fatigue (Spragg et al., 2022). In this section, we demonstrate that the power-law (a.k.a. Riegel) model implies a power–duration curve that naturally shifts downwards as the athlete becomes more fatigued without any additional modelling efforts.

**Figure 12:**
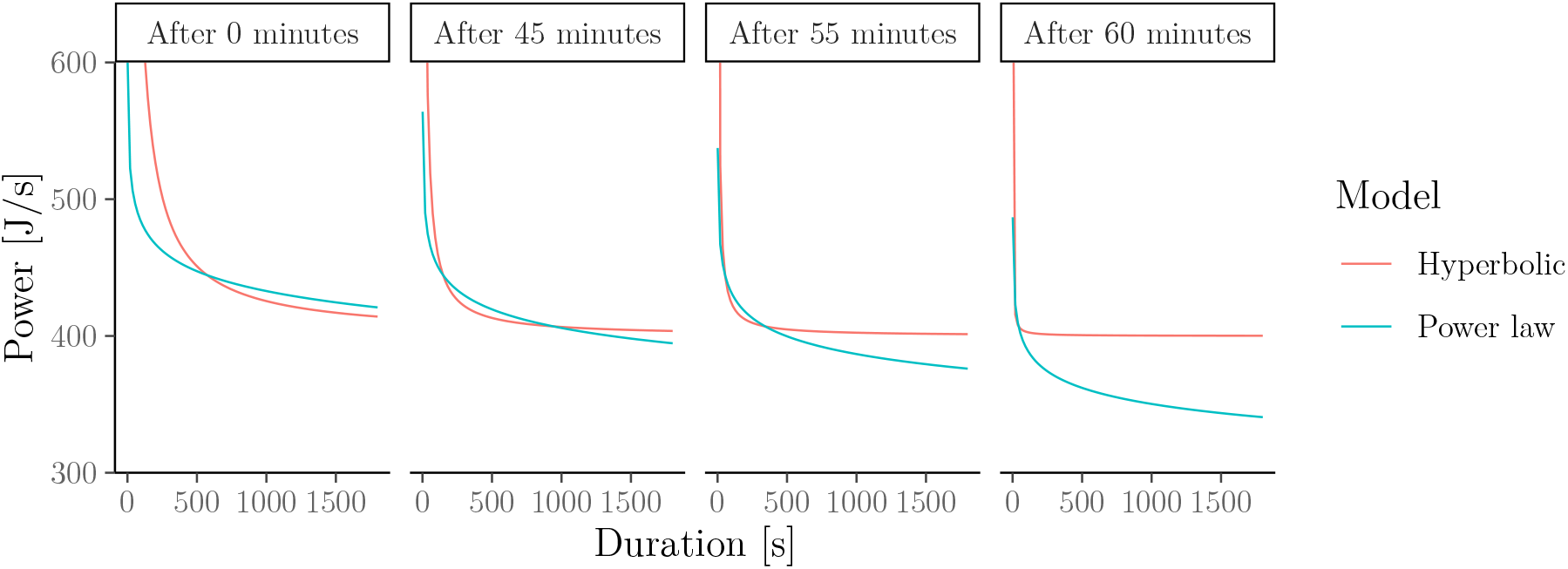
Fatigued power–duration relationship implied by the hyperbolic (a.k.a. critical-power) and power-law (a.k.a. Riegel) models for a cyclist who has already exercised 0, 45, 55 or 60 minutes at constant power output *P* = 407 J/s. The figure illustrates the downward shift of the power–duration curve under the power-law model (but not under the hyperbolic model) as the athlete becomes more fatigued.

### 5.2 Fatigued power–duration relationship

Assume that an athlete exercises at some constant intensity *P* > 0 for some duration *t* ≥ 0. If exhaustion has not yet set in at time *t*, then we may be interested in the power–duration relationship of this already partially fatigued athlete. We call this the *fatigued* power–duration relationship.

The specific form of the fatigued power–duration relationship depends on the chosen model for the fresh (i.e. non-fatigued) power–duration relationship. Figure 12 illustrates that the hyperbolic and power-law model imply a quite different behaviour of the fatigued power–duration relationship of a cyclist who has already exercised at power output *P* = 407 J/s for *t* = 0, 45, 55 or 50 minutes. The model parameters are *W*′ = 25 500, CP = 400, *F* = 1.05, and *S* is chosen such that the time to exhaustion at power output *P* is the same in both models. Figure 12 illustrates the following.

- **Hyperbolic model.** Under the hyperbolic model, the fatigued power–duration relationship again follows a hyperbolic model but with *W*′ replaced by

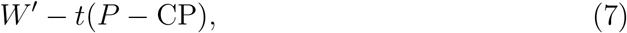

where *t* is in seconds.
- **Power-law model.** Under the power-law model, the fatigued power–duration relationship again follows a power-law model but with *S* replaced by

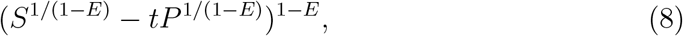

where *t* is in seconds.

The result in (7) follows directly from Section 2.2.4; the result in (8) is a consequence of the novel “rate-of-exertion interpretation” of the power-law model which we introduced in Section 2.3.4. More details are given in Appendix D.

### 5.3 Problems of the “critical power decreases with fatigue” hypothesis

The behaviour of the hyperbolic model shown in Figure 12 (i.e. the implication that an athlete can always maintain power output equal to at least CP even when heavily fatigued) contradicts empirical evidence and it has therefore been argued that CP and *W*′ should be treated as dynamic quantities that diminish with fatigue (Clark et al., 2019; Spragg et al., 2022). However, assuming that CP varies with exercise-induced fatigue has two drawbacks.

1. **Not inherent in the model.** This assumption requires additional modelling and data-collection efforts.
2. **Logical contradiction.** This assumption is also contradictory because if CP decreases with exercise (specifically: with exercise-induced fatigue) then the power–duration relationship is actually no longer hyperbolic (neither for a fresh nor for a fatigued athlete), i.e. the notion of a “rested” or “fatigued” critical power is not even well defined. To our knowledge, this problem has not been pointed out or addressed in the literature.

In contrast, for the power-law model, Figure 12 illustrates that the power–duration curve is naturally scaled downwards as the athlete becomes more fatigued. Crucially, this is inherent in the model, i.e. it does not lead to any logical contradictions and no additional data-collection or modelling efforts are needed.

## 6 Contribution IV: Relationship with popular performance predictors

### 6.1 Overview

In this section, we demonstrate that various other widely used performance predictors such as FTP (Allen and Coggan, 2012) in cycling, Jack Daniels’ “VDOT” calculator (Daniels and Gilbert, 1979) in running, Jeff Galloway’s “Magic Mile” (Galloway, 1984) in running, and Kurt Jensen’s “Golden Standard” (ergrowing.com, nd) in rowing can all be viewed as approximating a power-law model with a fixed endurance parameter *E*.

Hence, by the arguments from Section 2.3.3, the predictors discussed in this section are justifiable as a bias–variance trade-off when the available data points are too few (or too noisy) to accurately estimate *E*, e.g. when attempting to predict the finish time in a race from a single previous effort (note that at least two data points would be needed to estimate *E* in addition to *S*). However, when enough high-quality data are available, using the power-law model with not only *S* but also *E* estimated from the data will likely yield better predictions since *E* should be different for different individuals.

### 6.2 “Functional Threshold Power” (FTP) in cycling

In cycling, the so-called *functional threshold power* (*FTP*) is defined as the average power that a athlete can sustain for one hour. It is typically estimated by taking 95% of MMP_20_ (Allen and Coggan, 2012) or 90% of MMP_8_ (Carmichael and Rutberg, 2012), where MMP_*x*_ denotes the mean power output during an *x*-minute maximal effort.

FTP is the “de facto standard” for fitness assessment in cycling and used as a basis for prescribing training intensities by popular cycling analytics platforms such as Training Peaks, Garmin Connect and Zwift (Chorley and Lamb, 2020). Yet, the 95-% rule has been criticised

a. for predicting only a single point on the power–duration curve (Leo et al., 2022a),
b. for lacking a theoretical foundation (Mackey and Horner, 2021), and
c. for being inconsistent with the hyperbolic (a.k.a. critical-power) model:

> “determination of the FTP is based on the arbitrary subtraction of 5% of the mean power output during a 20-min [time trial]. The work performed above FTP was not equivalent to *W*′ and thus the physiological bases to the FTP protocol is unclear.” (Morgan et al., 2019)

We now show that Criticisms a, b and c are addressed (at least partly) if we take the view that the power–duration relationship follows a power law. That is, using the fatigue factor for average-aged male cyclists from Riegel (1981), i.e. *F* = 1/*E* = 1.04834 (see Table 1), the power-law model implies the “95-%” rule from Allen and Coggan (2012) because:

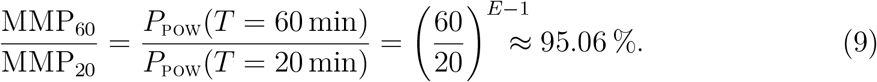

Figure 13 visualises the identity from Equation 9.

**Figure 13:**
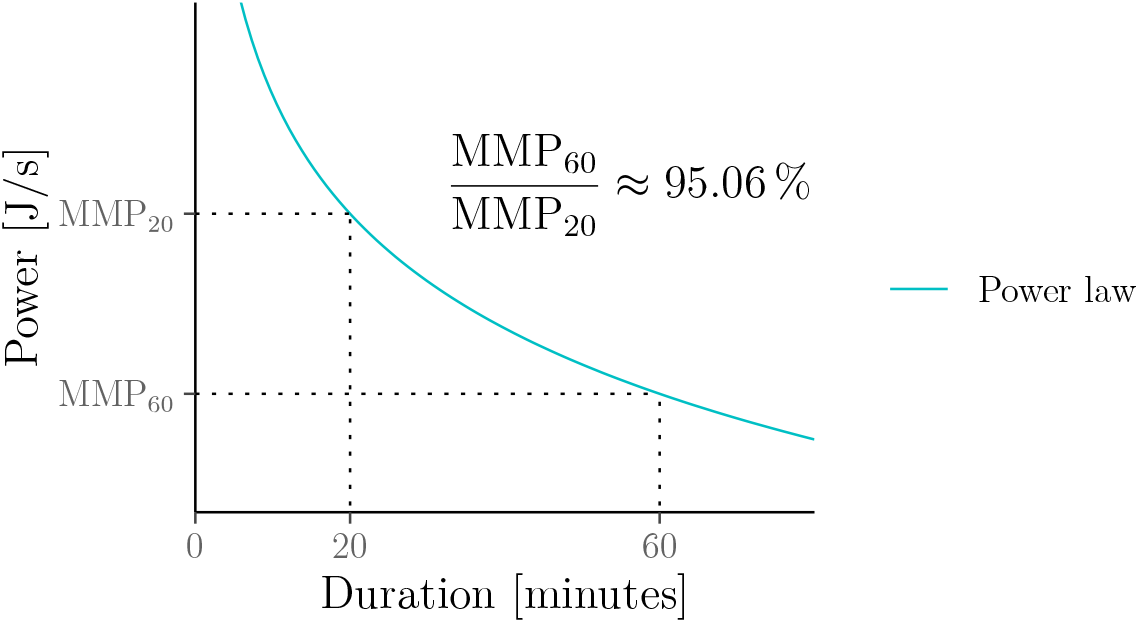
The connection between the 95-% rule for estimating FTP and the powerlaw model with endurance parameter from Riegel (1981); MMP_*x*_ denotes the (average) power output which the athlete can sustain over *x* minutes.

More generally, the power-law model implies that MMP_60_ can be estimated from an *x*-minute effort by taking the proportion (60/*x*)^*E*–1^ of MMP_*x*_, although larger durations *x* will likely yield more reliable estimates. The power-law model (again with *F* = 1/*E* = 1.04834) is thus also consistent with the “90-%” rule from Carmichael and Rutberg (2012) because (60/8)^*E*–1^ ≈ 91.12%. To our knowledge, this link between the rules for estimating FTP from 8-minute and 20-minute efforts and the power-law model has not been established in the literature.

We conclude by recalling that the fatigue factor *F* = 1.04834 derived by Riegel (1981) is based on world records across different athletes, i.e. it likely overestimates the endurance of most athletes. Whilst this may be somewhat counterbalanced by the fact that the FTP testing protocol asks cyclists to do a hard 5-minute effort before starting their 20-minute effort, this could explain why the 95-% rule often still overestimates MMP_60_ (Borszcz et al., 2018; Tramontin et al., 2022).

### 6.3 Jack Daniels’ “VDOT” calculator in running

The “world’s best running coach” (Burfoot, 2009), Jack Daniels, suggests a mathematical model under which a runner’s velocity–distance relationship depends on a single parameter, 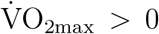. This parameter is typically unknown but can be estimated from a previous race result (this estimate is denoted as VDOT). Figure 14 shows that Jack Daniels’ predictor can again viewed as an approximation of the power-law model with fixed fatigue factor *F* = 1/*E* = 1.06 (which is well within the range of values for the fatigue factor given by Riegel (1981) for running and which is also the value used by the online finish-time predictor from Runner’s World Magazine (2013)).^12^ Of course, the predictions of both methods in Figure 14 do not coincide exactly. However, differences are likely to be negligible relative to the accuracy of estimating 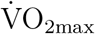. More details can be found in Appendix E.1.

**Figure 14:**
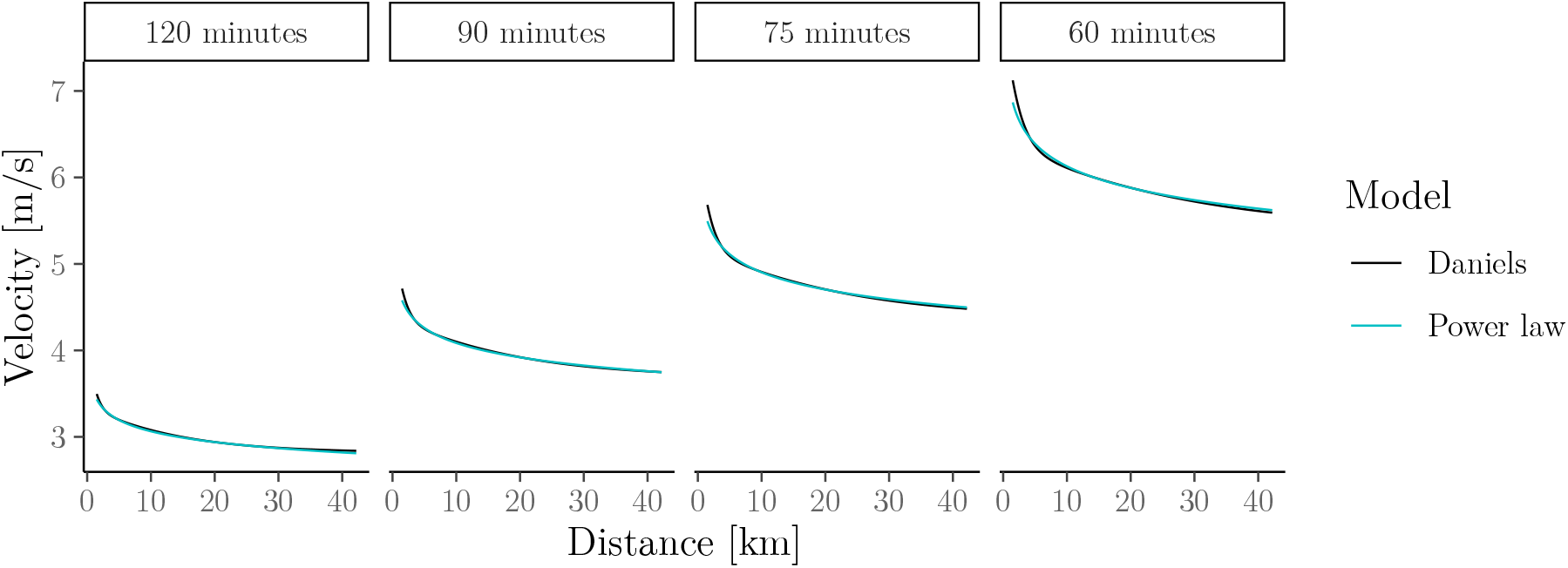
Velocity–distance relationship implied by Jack Daniels’ “VDOT” calculator compared with Riegel’s power-law model (with fixed fatigue factor *F* = 1.06) for athletes who can run a half marathon in 120, 90, 75, or 60 minutes, corresponding to VDOT values of around 36, 51, 63, and 81, respectively.

### 6.4 Jeff Galloway’s “Magic Mile” in running

Running coach and author Jeff Galloway suggests that the (average) pace achievable over 10-km, 10-mile, half-marathon and marathon distances can be estimated by multiplying the average pace achieved over one mile by 1.15, 1.175, 1.2 and 1.3, respectively (Galloway, 1984). Figure 15 illustrates that this rule can again be viewed as an approximation of the power-law model in which the fatigue factor is fixed to the value given for male runners in Riegel (1981), i.e. *F* = 1/*E* = 1.07732 (see Table 1).^13^ Of course, the predictions of both methods in Figure 15 do not coincide exactly. However, differences are likely to be negligible relative to the accuracy of estimating the mile pace. More details can be found in Appendix E.2.

**Figure 15:**
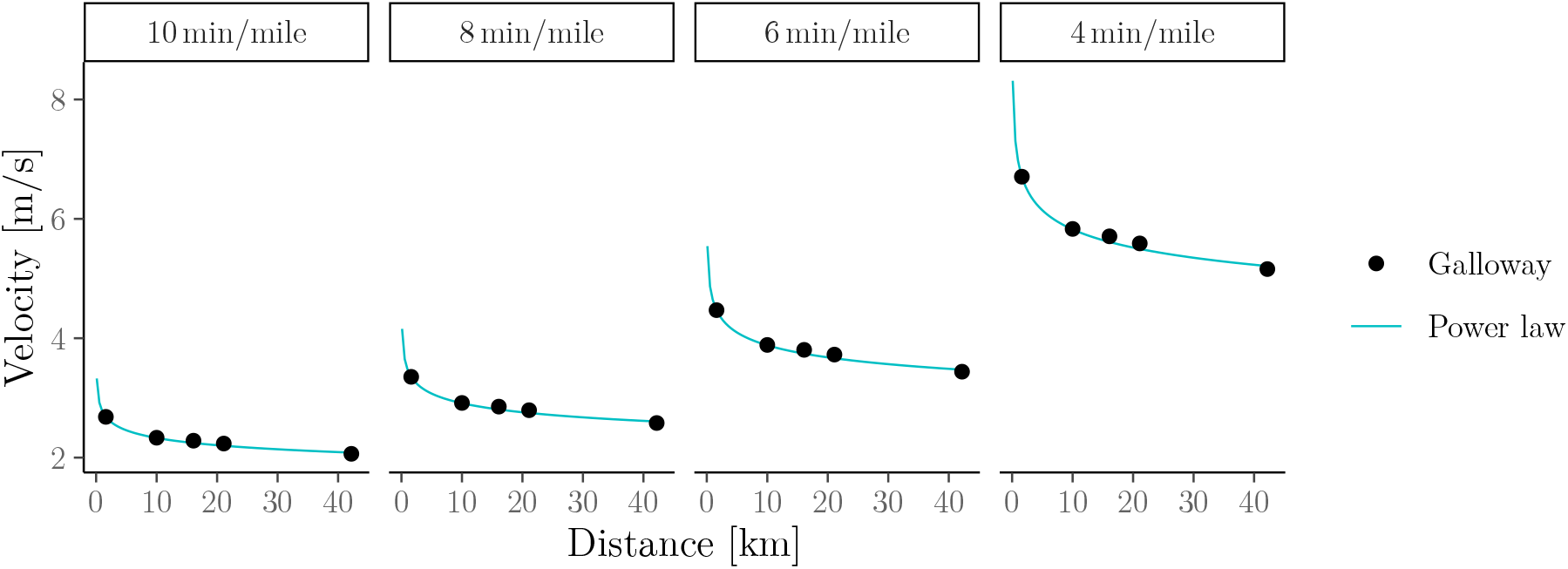
Velocity–distance relationship implied by Galloway’s “Magic Mile” and the power-law model (with fixed fatigue factor *F* = 1.07732 from Riegel 1981) for athletes that can run a mile in 10, 8, 6, or 4 minutes, respectively.

### 6.5 Kurt Jensen’s “Golden Standard” in rowing

Danish physiologist Kurt Jensen suggests that the (average) power achievable over 10 s, 60 s, 6km, and 60 min on a rowing ergometer can be estimated by multiplying the average power achieved on a rowing ergometer over 2km by 1.73, 1.53, 0.85 and 0.76, respectively (ergrowing.com, nd; worldofrowing.com, 2019). Figure 16 illustrates that this rule can again be viewed as an approximation of the power-law model with fixed endurance parameter *E* = 0.85 (i.e. with fatigue factor *F* = 1/*E* ≈ 1.07732).^14^ Of course, the predictions of both methods in Figure 16 do not coincide exactly. However, differences are likely to be negligible relative to the accuracy of estimating the power output over 2km. More details can be found in Appendix E.3.

**Figure 16:**
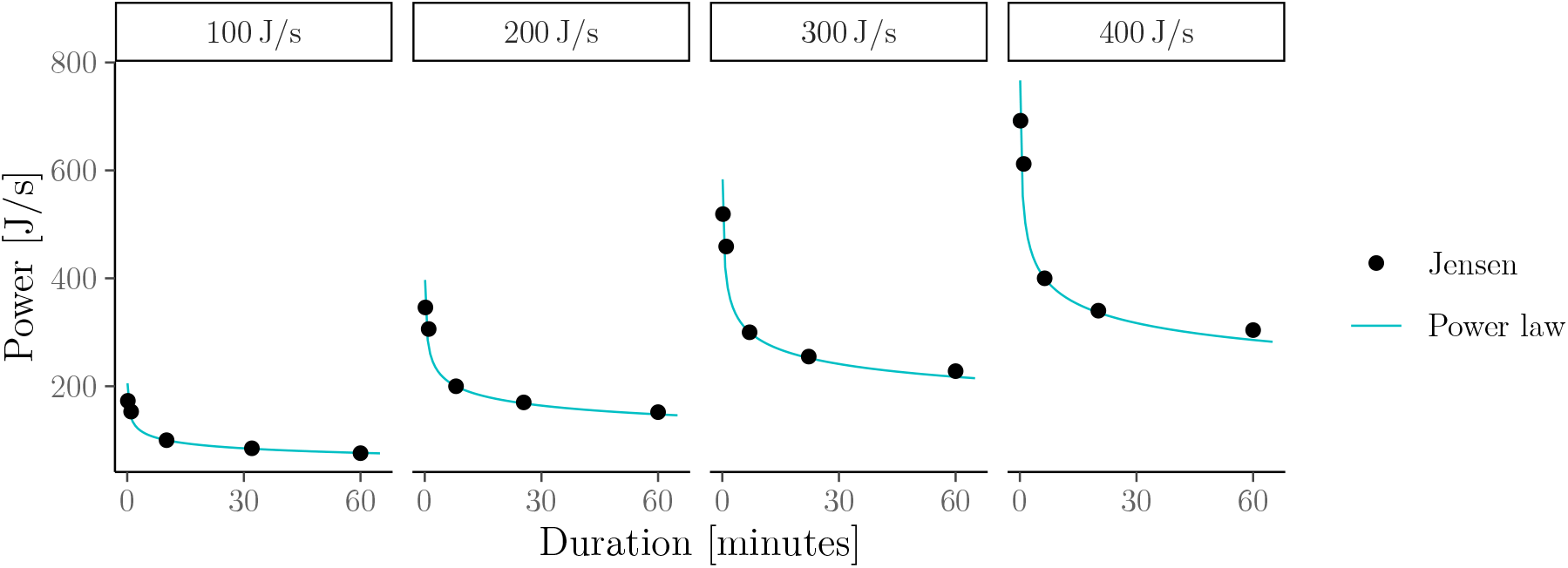
Power–duration relationship implied by Jensen’s “Golden Stadard” and the power-law model (with fixed endurance parameter *E* = 0.85) for athletes that can sustain 100, 200, 300, and 400 J/s, respectively, over 2km.

**Figure 17:**
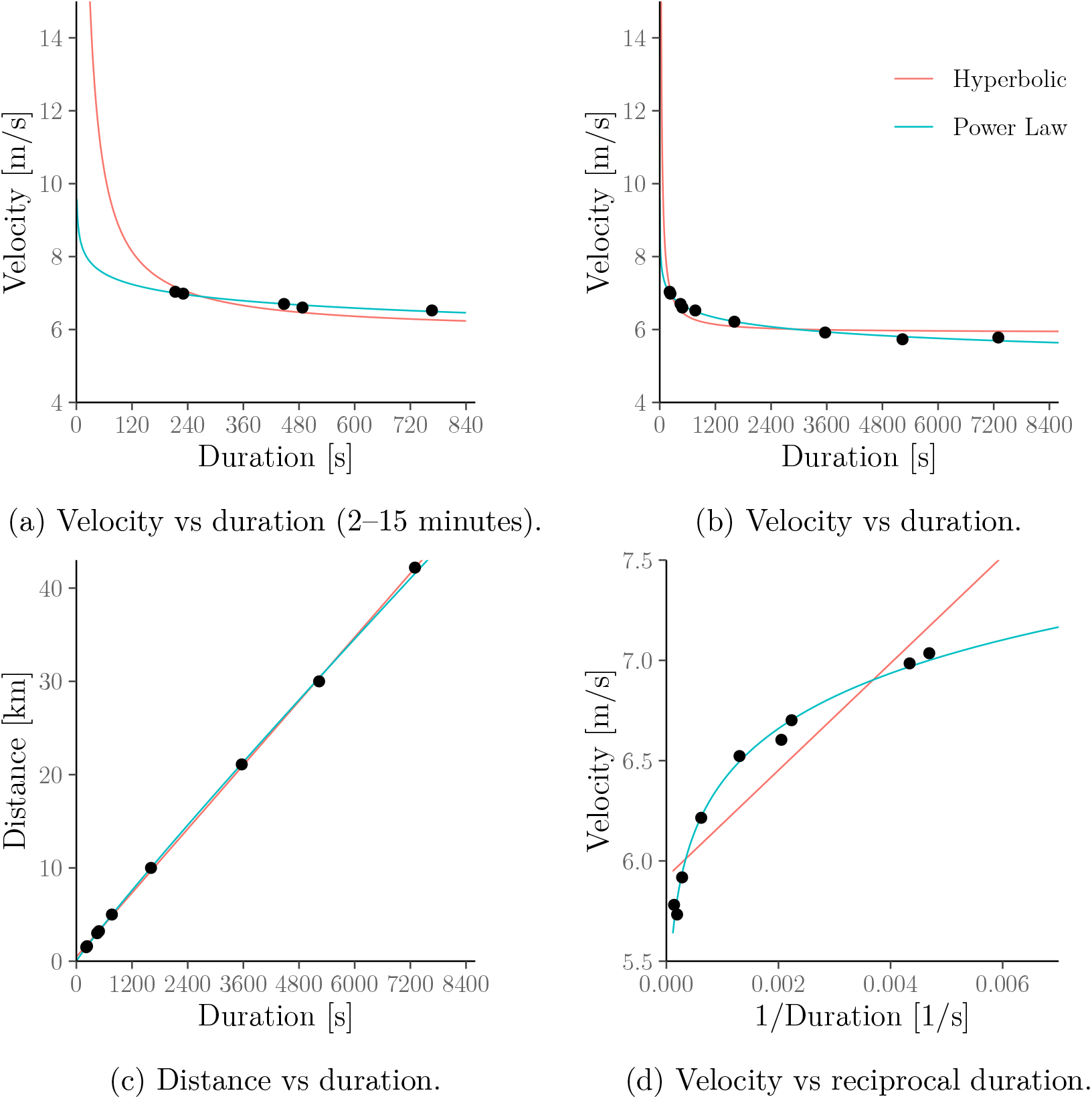
A reproduction of Figure 4 but the hyperbolic (a.k.a. critical power) model is fitted to all available data. This reduces the error for shorter and longer durations but increases the error in the 2–15 minute range.

**Figure 18:**
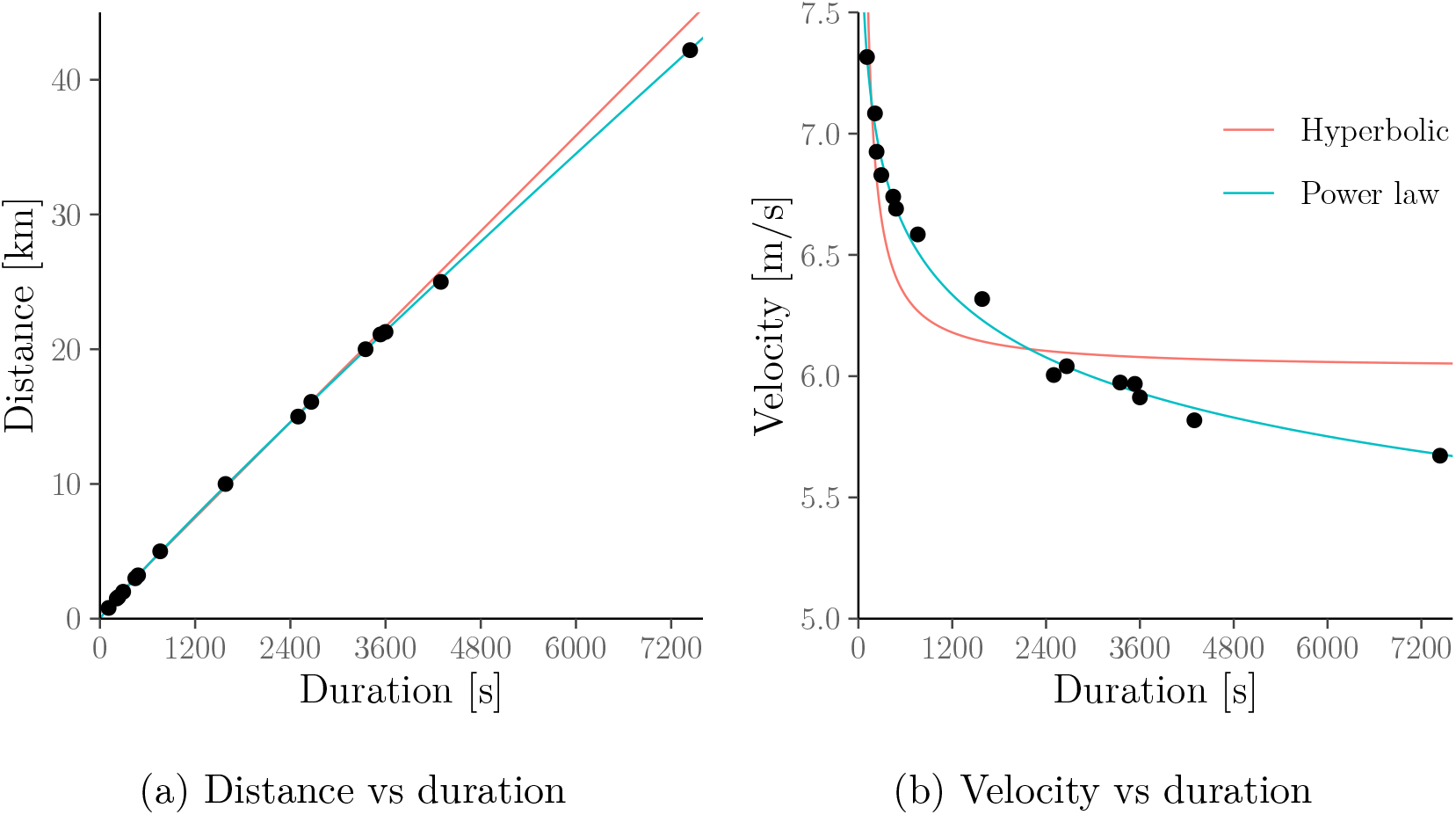
A reproduction of Figure 5 but the hyperbolic (a.k.a. critical power) model is fitted to all available data. This reduces the error for shorter and longer durations but increases the error in the 1500–15 000m range.

**Figure 19:**
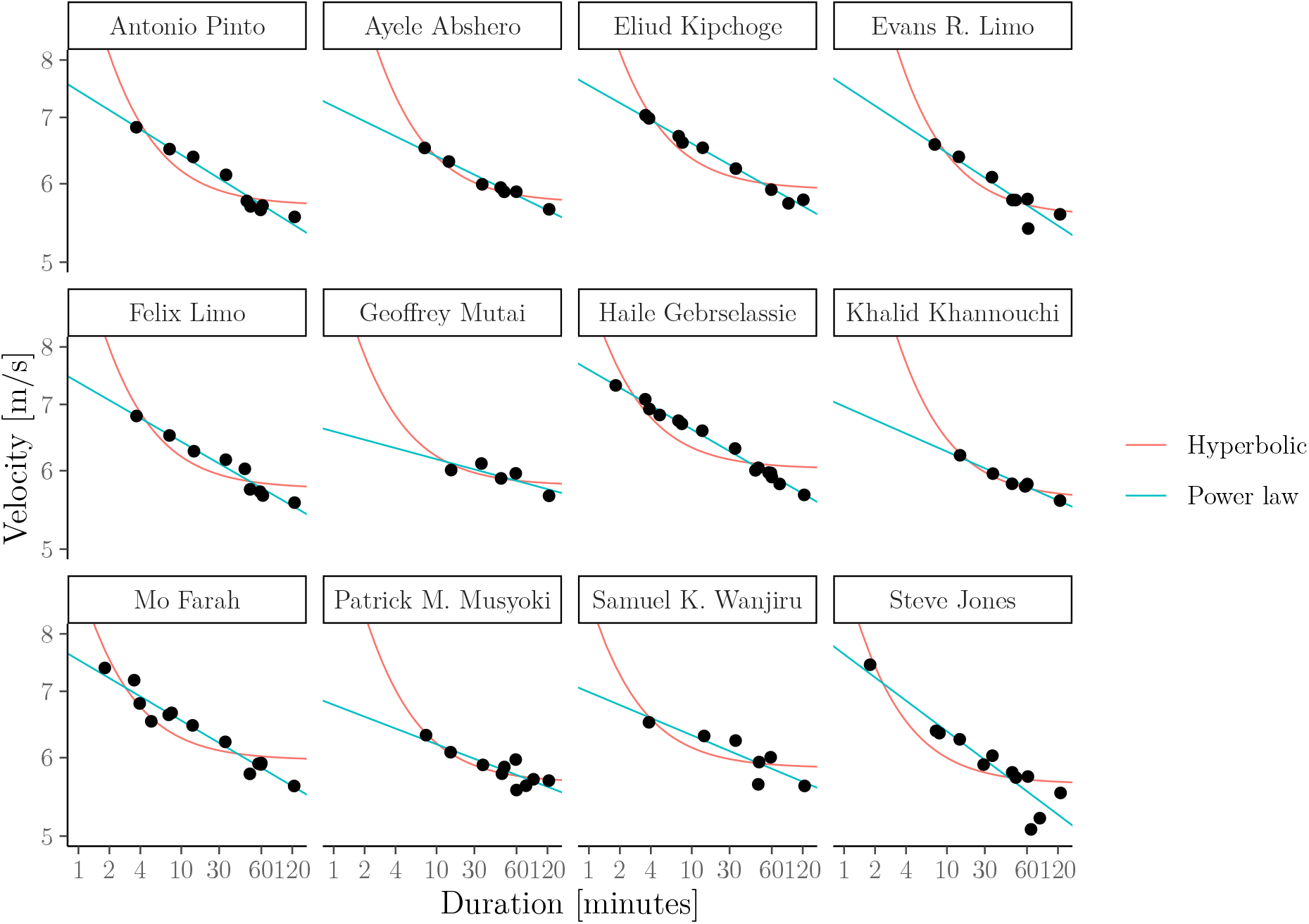
A reproduction of Figure 6 but the hyperbolic (a.k.a. critical power) model is fitted to all available data. This reduces the error for shorter and longer durations but increases the error in the 1500–15 000m range.

## 7 Discussion

### Summary

We have demonstrated – both empirically and theoretically – that the power–duration relationship is more adequately represented by the power-law (a.k.a. Riegel) model than by the hyperbolic (a.k.a. critical-power) model. In particular, our work highlights the following.

1. The power-law model fits power–duration data, e.g., in cycling or rowing, and race results in running better than the hyperbolic model over a wide range of exercise durations or race distances.
2. The power-law model is applicable to a wide range of exercise intensities, durations or distances. In contrast, the hyperbolic is restricted to a very limited range of durations, i.e. between 2 and around 15 to 25 minutes.
3. The power-law model only has two parameters, just like the hyperbolic model, and is just as easy to fit.
4. The power-law model appears to be a safer tool for pace selection than the hyperbolic model (except in short sprint races) because it accounts for the fact that overpacing, e.g. sprinting off on the first kilometre of a long-distance race, is detrimental to the overall performance.
5. The power-law model implies that the power–duration relationship changes with fatigue in a manner that is in closer agreement with empirical evidence than the fatigued power–duration relationship implied by the hyperbolic model;
6. The power–law model (but not the hyperbolic model) is consistent with many rules-of-thumb for performance prediction which are popular among practitioners such as FTP in cycling, Jack Daniels’ “VDOT” calculator and Jeff Galloway’s “Magic Mile” in running, or Kurt Jensen’s “Golden Standard” in rowing.

We stress that we do not claim that the power-law model is “correct”. Indeed, both models likely constitute gross simplifications of reality. For instance, Blythe and Király 2016 found that it can be improved by including additional (individual) correction factors.

However, given the choice between the hyperbolic model and the power-law model, Points 1-6 above leave very little reason for choosing the former over the latter for fitness assessment or performance prediction in athletes. The only exception is the modelling of intermittent exercise because, as discussed in the “Limitations” below, the power-law model cannot easily incorporate recovery.

### Implications for the notion of “critical power”

It is important to recognise that the term “critical power” (similarly: “critical velocity” or “critical speed”) can refer to two notions which are, in principle, distinct:

**CP_asymp_** The power asymptote of the hyperbolic model, i.e. the parameter CP.
**CP_thresh_** A threshold which is thought to separate distinct physiological responses, and by extension, the “heavy” from the “severe” exercise intensity domain.

Note that critical power in the sense of **CP_asymp_** only exists because the power–duration relationship is modelled by a hyperbolic function – it does not exist if the data are modelled via a power law which, our work suggests, is the more appropriate functional form. In this precise sense, critical power could be called a “mathematical artefact” as is frequently done by its critics (e.g., Gorostiaga et al., 2022b). The same applies to finite work capacity above critical power, *W*′. Athletes and coaches should keep this in mind when assessing the meaningfulness of CP and *W*′ as metrics for fitness assessment.

We stress that the previous interpretation only concerns **CP_asymp_**. **CP_thresh_**, i.e. the notion of critical power as a “physiological threshold” is, in principle, a completely separate concept. Nonetheless, despite being a physiological threshold, **CP_thresh_** is not easily determined from physiological measurements. Instead, it is commonly argued that **CP_thresh_** coincides with **CP_asymp_** so that it can be identified by fitting the hyperbolic model to power–duration measurements. Implicit in this practice is the assumption that the changes in physiological responses which separate the “heavy” from the “severe” exercise intensity domain manifest themselves in a “levelling off” of the power–duration curve towards **CP_thresh_** as durations increase towards, say, 15 or 20 minutes. However, our work suggests that this is not the case. Indeed, if it was the case, the hyperbolic model would fit better than the power-law model for durations up to 15 or 20 minutes.

### Implications for training prescription

Athletes and coaches often use critical power/velocity (in the sense of **CP_asymp_**) to select training intensities. We stress that setting training intensities is still possible if we instead use the power-law model. In fact, this is consistent with existing practice. For instance,

- in cycling, training intensities are often based on the power that a athlete can sustain for some specific duration (e.g., 60 minutes in the case of FTP);
- in running, training velocities are often based on the velocity that the runner can sustain over some specific distance (e.g., “5-km pace”, “marathon pace” or “half-marathon pace”).

In both cases, the power-law model gives a straightforward and principled way of estimating the required powers/velocities, e.g. it allows athletes to easily calculate how fast an individual athlete’s “marathon pace” actually is.

### Limitations

The power-law model also has two limitations (though the first is shared by the hyperbolic model):

1. The power-law model assumes that athletes have to come to a complete stop from one second to the next when reaching the limit implied by the power–duration curve (i.e. when their “exertion” in (6) reaches 1). For instance, the power-law model cannot explain the gradual decrease in power output observed in a 3-minute all-out test (Burnley et al., 2006, Figure 3A) (however, neither can the hyperbolic model). This property may also mean that the penalty for overpacing shown in Figure 11 is too harsh.
2. Unlike the hyperbolic model, the power-law model cannot easily be extended to incorporate recovery (such as in Morton and Billat 2004) due to lack of a “critical power” threshold below which recovery can be assumed to occur. We could, of course, easily add a positive constant to (3) as in Tsai (2015) to obtain such a threshold but this would again imply that there exists a non-zero power output which can be sustained for a “very long time without fatigue”.

We are currently developing a new model which retains the advantages of the power-law model whilst resolving both Limitations 1 and 2 (Drake et al., nd).

## Acknowledgements

We are very grateful to Dr Mark Burnley for valuable comments on an initial version of the manuscript. We would like to thank Dr Peter Leo for providing the power–duration data used in Figure 10.

## A Additional details for Section 2

### A.1 Hyperbolic (a.k.a. critical-power) model

#### A.1.1 Equivalent relationships

Using simple linear algebra, we can verify that the hyperbolic model implies the following relationships which are also illustrated in Figure 1. For *W*′ > 0, *P* > CP ≥ 0 and *W* = *PT* > *W*′:

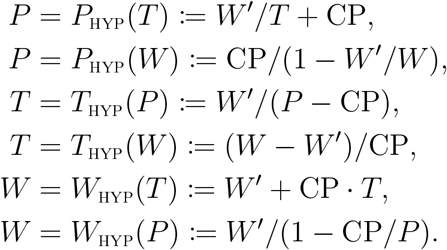

As discussed in Section 2.1, we sometimes replace “power” (*P*) by “velocity” (*V*) and “work” (*W*) by “distance” (*D*). In this case, we also write *V*_HYP_ and *D*_HYP_ instead of *P*_HYP_ and *W*_HYP_.

### A.2 Power-law (a.k.a. Riegel) model

#### A.2.1 Equivalent relationships

Using simple linear algebra, we can again verify that the power-law model implies the following relationships which are also illustrated in Figure 2. For *S* > 0, *F* = 1/*E* > 1 and *W* = *P* · *T* > 0:

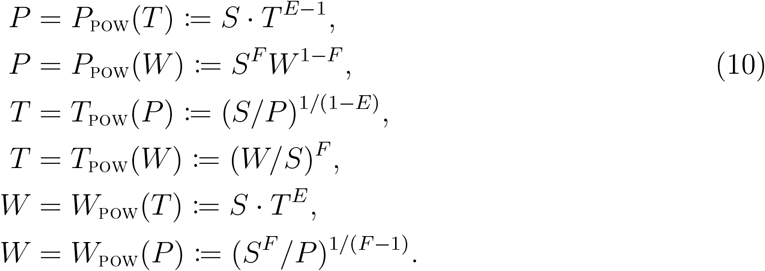

As discussed in Section 2.1, we sometimes replace “power” (*P*) by “velocity” (*V*) and “work” (*W*) by “distance” (*D*). In this case, we also write *V*_POW_ and *D*_POW_ instead of *P*_POW_ and *W*_POW_.

#### A.2.2 Bias–variance trade-off and Riegel-predictors

Recall that, as mentioned in Section 2.3.3, sufficiently many data points are needed to accurately estimate the speed parameter, *S*, and endurance parameter, *E*, (equivalently: the fatigue factor *F* = 1/*E*) of the power-law model.

In practice, there is often a need to form predictions based on a very small number of data points – possibly just a single observation, e.g. when predicting a marathon finish time from a single prior half-marathon result.

In this case, it is common to fix *E* (equivalently: *F* = 1/*E*) to a suitable default value which then permits the estimation of *S* from a single data point. Assume that we have a previous power measurement *P*_0_ recorded over some duration *T*_0_. Then solving (3) for *S* and using (*P*_0_, *T*_0_) in place of (*P, T*), gives the estimate 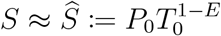; and plugging 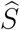 back into (3) implies the power–duration relationship:

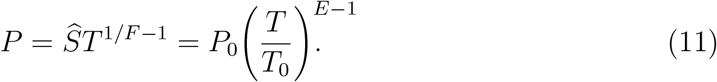

When applied to, e.g., running, this gives an easy way of predicting the finish time *T* in a race over distance *D* from the finish time *T*_0_ in a previous race over some other distance *D*_0_. More specifically, after replacing power *P* by velocity *V* = *D/T* (see Section 2.1) and *E* by 1/*F*, we can re-arrange (11) as

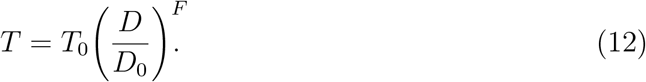

For instance, the finish-time calculator from Runner’s World Magazine (2013) implements (12) with *F* = 1.06 for all runners. This comes with the assumption that the user is an average runner with without “a natural significant bias towards either speed or endurance” (Runner’s World Magazine, 2013).

Of course, the fatigue fatigue factor *F* (equivalently: the endurance parameter *E* = 1/*F*) is likely different for each person as shown in Blythe and Király (2016); Zinoubi et al. (2017). Hence, setting *F* (equivalently: *E*) to a default value incurs a bias. However, this bias is outweighed by the variance reductions brought about by avoiding the need for estimating *F* from limited amounts of data. Thus, fixing the fatigue factor to a sensible default value can be justified as a *bias–variance trade-off*.

## B Additional details for Section 3

### B.1 Model behaviour over short durations

The hyperbolic (a.k.a. critical-power) model implies that any sufficiently small amount of work 0 < *W* < *W*′ can be generated arbitrarily quickly. For instance, if the model was applied to short durations, it would thus imply that an elite runner can “teleport” over more than one hundred metres. In contrast, power-law (a.k.a. Riegel) model has no such unrealistic implication. Informally, these results can be seen from Figures 1c and 2c which show that the second-axis intercept of the work–duration curve is

- *W*′ > 0 under the hyperbolic model;
- 0 under the power-law model.

More formal proofs – which are needed because the power–duration relationship *P*(*T*) is not actually well defined at *T* = 0 – are given in Propositions 3 and 4.

#### Proposition 3.

*Let* 0 < *W* < *W*′. *Then for any T* > 0, *W* < *T* · *P_HYP_*(*T*).

**Proof.** Let *T* > 0. Then since *W* < *W*′,

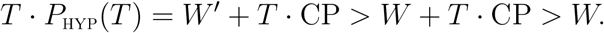

This completes the proof.

#### Proposition 4.

*Let S* > 0 *and F* > 1. *Then for any W* > 0, *there exists a unique duration T* > 0 *such that W* = *T* · *P_POW_*(*T*).

**Proof.** The function 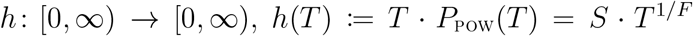 is strictly increasing and *h*(0) = 0. Thus, there exists a unique value *T* > 0 such that *W* = *h*(*T*) = *T* · *P*_POW_(*T*).

### B.2 Case study from Jones et al. (2019)

Table 2 shows Eliud Kipchoge’s personal records in different running events (as of 5 November 2021). The data were collected from World Athletics (2021). Note that the difference between, e.g., the 5000m and 5 km event is that the former was run on the track whereas the latter was run on the road.

**Table 2:**
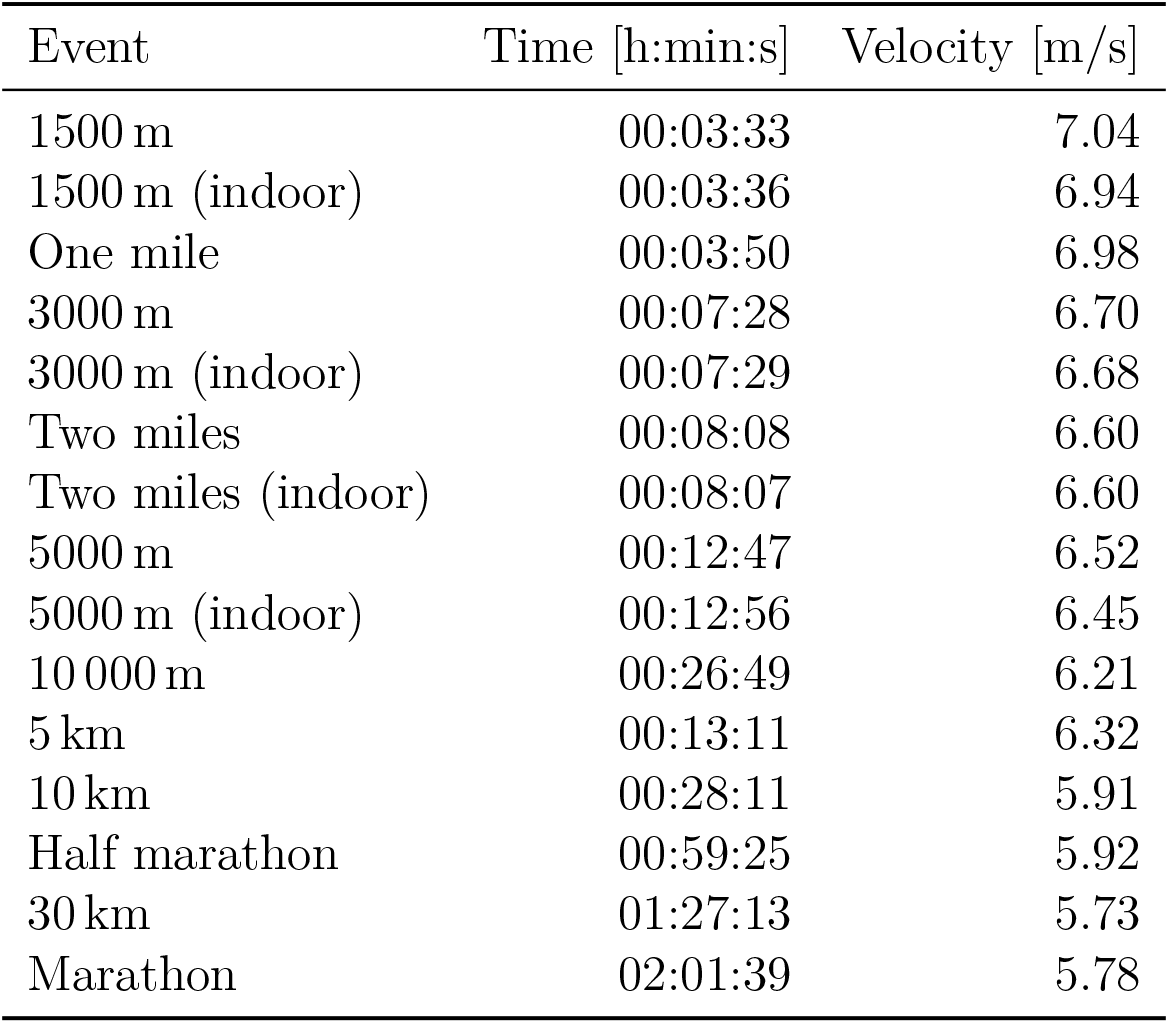
Personal records of Eliud Kipchoge in different events.

We now provide a version of Figure 4 in which we fit the hyperbolic model to all personal records up to marathon distance in the same way as the power-law model.

### B.3 Case study from Jones and Vanhatalo (2017)

Table 3 shows Haile Gebrselassie’s personal records in different running events. The data were collected from World Athletics (2021). Note that the difference between, e.g., the 10 000m and 10 km event is that the former was run on the track whereas the latter was run on the road.

**Table 3:**
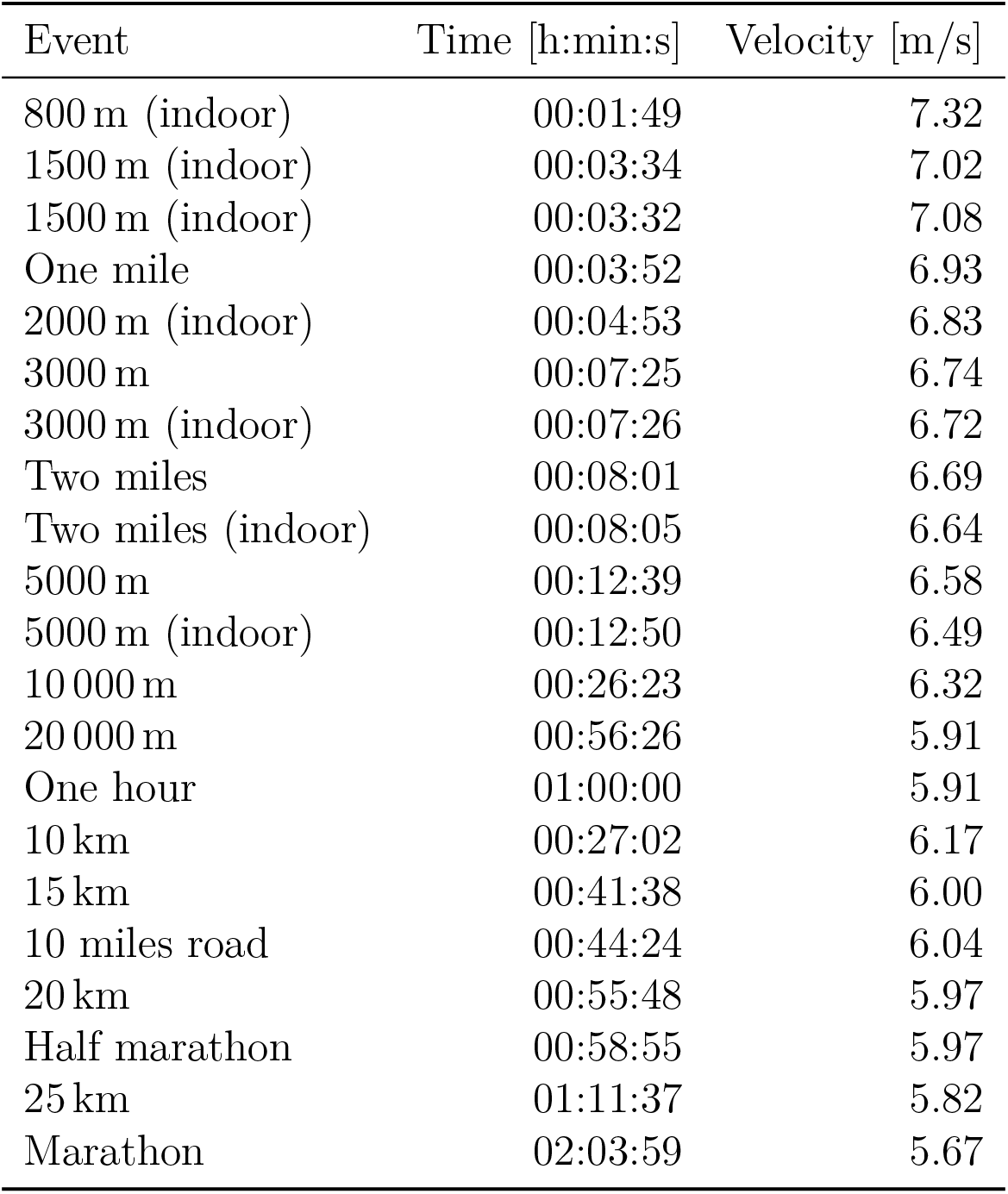
Personal records of Haile Gebrselassie in different events.

| Event | Time [h:min:s] | Velocity [m/s] |
| --- | --- | --- |
| 800 m (indoor) | 00:01:49 | 7.32 |
| 1500 m (indoor) | 00:03:34 | 7.02 |
| 1500 m (indoor) | 00:03:32 | 7.08 |
| One mile | 00:03:52 | 6.93 |
| 2000 m (indoor) | 00:04:53 | 6.83 |
| 3000 m | 00:07:25 | 6.74 |
| 3000 m (indoor) | 00:07:26 | 6.72 |
| Two miles | 00:08:01 | 6.69 |
| Two miles (indoor) | 00:08:05 | 6.64 |
| 5000 m | 00:12:39 | 6.58 |
| 5000 m (indoor) | 00:12:50 | 6.49 |
| 10 000 m | 00:26:23 | 6.32 |
| 20 000 m | 00:56:26 | 5.91 |
| One hour | 01:00:00 | 5.91 |
| 10 km | 00:27:02 | 6.17 |
| 15 km | 00:41:38 | 6.00 |
| 10 miles road | 00:44:24 | 6.04 |
| 20 km | 00:55:48 | 5.97 |
| Half marathon | 00:58:55 | 5.97 |
| 25 km | 01:11:37 | 5.82 |
| Marathon | 02:03:59 | 5.67 |

We now provide a version of Figure 5 in which we fit the hyperbolic model to all personal records up to marathon distance in the same way as the power-law model.

We now provide a version of Figure 6 in which we fit the hyperbolic model to all personal records up to marathon distance in the same way as the power-law model.

Finally, we provide a numerical comparison of the relative errors of both models for the athletes used in Section 3.3.

### B.4 Model-error computation

Here, we explain more formally how the model errors in Sections 3.4, 3.5, and 3.6 and in Table 4 from Appendix B.3 are calculated.

**Table 4:** Relative errors of the hyperbolic (a.k.a. critical-power) and power-law (a.k.a. Riegel) models for the 12 athletes from Jones and Vanhatalo (2017) fitted either to personal records over 1500–15 000m or all available personal records (up to marathon distance). Errors are calculated as the relative distance between observed and predicted velocities as explained in Appendix B.4. Colours highlight the better-performing model.

| Athlete | 1500–15 000 m |  | 0–42 195 m |  |
| --- | --- | --- | --- | --- |
|  | Hyperbolic | Power law | Hyperbolic | Power law |
| Antonio Pinto | 2.1700 | **1.0000** | 2.6900 | **1.1200** |
| Ayele Abshero | 0.5700 | **0.4700** | 0.9700 | **0.5600** |
| Eliud Kipchoge | 0.8400 | **0.3500** | 2.5000 | **0.6000** |
| Evans Rutto Limo | 1.6600 | **0.7600** | 2.0300 | **1.6900** |
| Felix Limo | 0.7500 | **0.5600** | 2.8100 | **1.1300** |
| Geoffrey Mutai | 1.2200 | **1.1800** | 1.7100 | **1.3900** |
| Haile Gebrselassie | 1.5100 | **0.7900** | 3.2300 | **0.5700** |
| Khalid Khannouchi | 0.2900 | **0.0400** | 0.6800 | **0.4400** |
| Mo Farah | 1.7200 | **1.4300** | 3.1500 | **1.4500** |
| Patrick Makau Musyoki | **0.3400** | 0.3500 | **1.2900** | 1.5000 |
| Samuel Kamau Wanjiru | 3.3900 | **2.6500** | 3.0200 | **1.9800** |
| Steve Jones | 1.1200 | **0.7500** | 4.0900 | **2.8400** |
| Average | 1.2983 | **0.8608** | 2.3417 | **1.2667** |

Assume that (after cleaning), our data set contains *N* athletes and that *K_n_* ≥ 2 pairs of power and duration (or velocity and duration, in the case of running) measurements 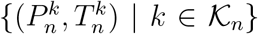 are available for the *n*th athlete. Then the average relative error of the hyperbolic model for durations between *T*_0_ and *T*_1_ is calculated as follows.

1. Let 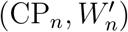 be the estimate of (CP, *W*′) obtained via linear regression using only data corresponding to activities whose duration is no shorter than *T*_0_ and no longer than *T*_1_, i.e. using only the measurements 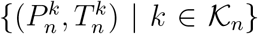, where 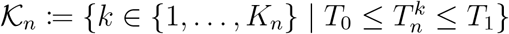.
2. Let 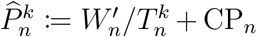 be the fitted power for the *k*th activity of the *n*th athlete under the hyperbolic model. The relative error of the hyperbolic model for the data from the *n*th athlete in this exercise duration range is then given by

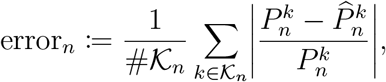

where 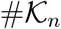 is the cardinality of 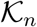 (i.e. the number of power–duration measurements for the *n*th athlete in this exercise duration range).
3. Then the average relative error of the hyperbolic model for this exercise duration range (i.e. the height of one of the red bars in Figures 7, 8 or 9) is then given by the following weighted average (where we weight by the number of available data points for each athlete):

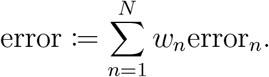

where we have defined the *n*th weight as:

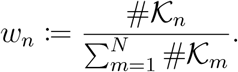
4. The associated standard error (i.e. the size of the error bar in Figures 7, 8 or 9) is then calculated as:

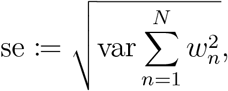

where

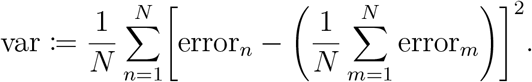 That is, if all weights are equal to 1/*N*, then 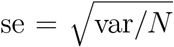 reduces to the usual standard error of the (unweighted) mean.

For the power-law model, we proceed analogously. The only difference is that we instead compute linear-regression estimates (*S_n_, E_n_*) of (*S, E*) and then define the fitted values as 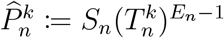.

### B.5 Large-data study in running

Here, we give additional details about the data set used for the study in Section 3.4. The data were collected by Blythe and Király (2016) and are available for download here: https://figshare.com/articles/dataset/thepowerof10/3408202. For the study, we removed all runners who have at most three race results with finishing times in the 2–15 minute range or at most two race results with finishing times in the 2–15 minute range over distinct distances.

### B.6 Large-data study in rowing

Here, we give additional details about the data set used for the study in Section 3.5. The data were collected on 28 August 2022 from www.nonathlon.com for the years (i.e. seasons) from 2002 to 2022. The data are given either as finish times *T* over fixed distances *D* (500m, 1km, 5km, 6km, 10km, 21.0975km and 42.195km) or as covered distances *D* over fixed durations (30 min and 60 min). To calculate the power output *P*, we use the commonly used conversion, where *T* is in seconds, *D* is in metres and *P* is in joules per second:

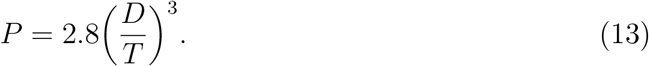

Finally, we removed all athlete seasons which have less than three records in the 2–20 minute range, and any athlete season which reported an (equivalent) average power output of over 1300 J/s leaving 3244 unique athlete seasons.

### B.7 Large-data study in cycling

We downloaded all the possible data (as of 9 June 2022) from Golden Cheetah using the GoldenCheetahOpenData (v0.2; Kosmidis, 2022) package in R (v4.1.2; R Core Team, 2021). We created a data set consisting of, for each athlete *n*, the highest measured average power outputs, MMP_*n,x*_, over given durations, *x* > 0. These are often referred to as *mean maximal power* (*MMP*) values. Before fitting the models, we excluded some data points. We now give more details.

- Power-meter malfunctions are not uncommon and can overestimate the power being produced by an athlete. To alleviate this problem, we removed rides which would have led to unrealistically high MMP values. This is done as follows, where *N* is the number of athletes and where *x* takes values in a set of 54 durations *x*, ranging from five seconds to two hours.
  1. Calculate “auxiliary” MMP values 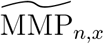 for each athlete *n* and duration *x* based on the original data set.
  2. Calculate the 95th percentile *q_x_* of 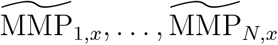.
  3. Discard any ride of any athlete *n* which contains an auxiliary MMP value which is such that 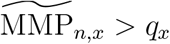.
  4. For each athlete *n* and each duration *x*, compute the MMP values MMP_*n,x*_ based on the remaining rides.
- Many of the MMP values calculated as above likely correspond to submaximal efforts which should not be used to estimate the models. Unfortunately, it is unknowable for which of these values this is the case. However, values MMP_*n,x*_ such that MMP_*n,x*_ ≤ MMP_*n,y*_, for some *x* < *y, cannot* be maximal and were therefore excluded. We stress that this does not necessarily mean that all the remaining data points correspond to maximal efforts.
- We removed athletes with fewer than three MMP values in the 2–15 minute range. This is because both models would have zero error if fitted to only two data points.

### B.8 Piecewise-defined models

In this section, we provide details on the “piecewise-defined” models from Péronnet and Thibault (1989); Puchowicz et al. (2020); Luttikholt and Jones (2022). Throughout, *T*_*_ > 0 denotes the threshold which is such that the power–duration relationship *P*(*T*) is (approximately) hyperbolic for durations *T* ≤ *T*_*_ whilst a different functional form is used for durations *T* > *T_*_*. Furthermore, we let 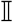 denote the indicator function, i.e. 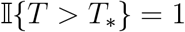 if *T* > *T*_*_ and 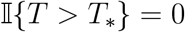, otherwise.

#### B.8.1 Péronnet and Thibault (1989)

Let *T*_*_ = 420 s (7 min) be the threshold parameter (denoted *T*_MAP_ in Péronnet and Thibault 1989) and let *k*_1_ = 30, *k*_2_ = 20, *f* = −0.233, and BMR = 1.2 be known constants. Furthermore, let *A*, MAP > 0 and *E* < 0 be other unknown (and athlete-specific) model parameters. Then the model from Péronnet and Thibault (1989) is

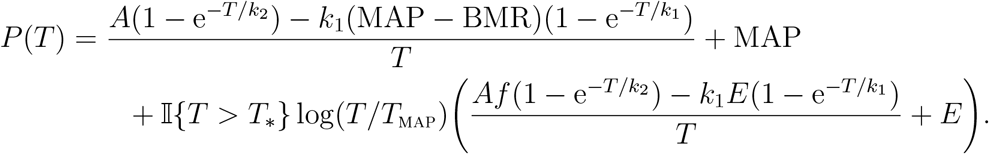

Figure 20 illustrates the model and shows that it has a “kink” in the power–duration curve at duration *T* = *T*_*_. Note that lim_*T*→∞_ *P*(*T*) = −∞.

**Figure 20:**
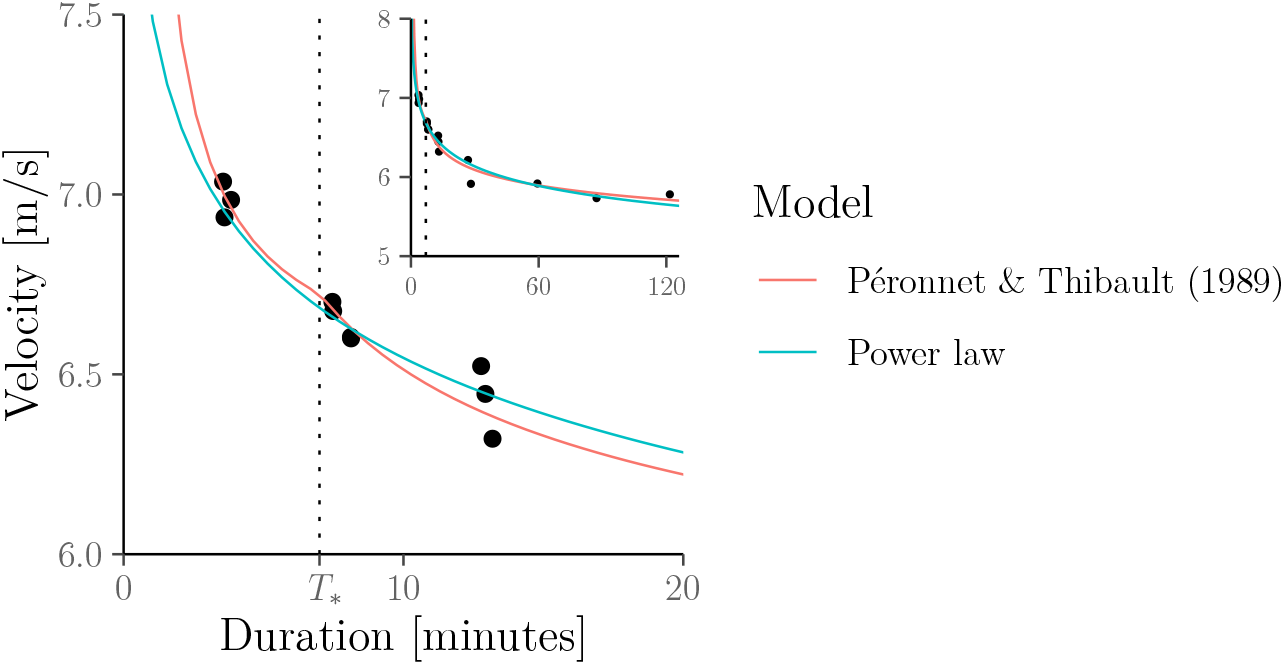
The velocity–duration relationship posited by the model from Péronnet and Thibault (1989) fitted to Eliud Kipchoge’s personal records.

#### B.8.2 Puchowicz et al. (2020)

Let *T*_*_ = 1800 s (30 min) be the threshold parameter (denoted TCPmax in Puchowicz et al. 2020) and let *W*′, CP, *P*_MAX_, *A* > 0 be unknown athlete-specific parameters. Puchowicz et al. (2020) propose the following three models.

- The *OmPD* model is given by

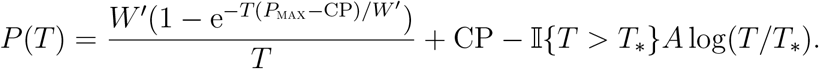
- The *Om3CP* model is given by

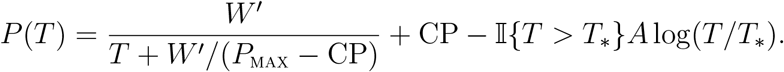
- The *OmExp* model is given by

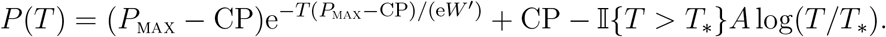

Figure 10 in the main manuscript illustrates these models and shows that they again have a “kink” in the power–duration curve at duration *T* = *T*_*_. Additionally, for all of these models, lim_*T*→∞_ *P*(*T*) = −∞.

#### B.8.3 Luttikholt and Jones (2022)

Let *T*_*_ = 360 s (6 min) be the threshold parameter and let *W*′, CP > 0 and 0 < *E* < 1 be athlete-specific parameters. Then the model from Luttikholt and Jones (2022) uses the hyperbolic (a.k.a. critical-power) model for durations up to *T*_*_ and uses a power-law model for durations longer than *T*_*_, i.e.:

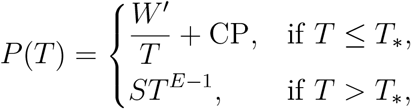

where continuity of the power–duration curve is ensured by setting

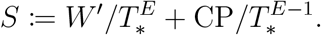

Figure 21 illustrates the model and shows that it again has a “kink” in the power–duration curve at duration *T* = *T*_*_. However, this kink is not very pronounced due to the small value of the threshold *T*_*_, i.e. because the hyperbolic model is only used for durations up to 6 minutes here.

**Figure 21:**
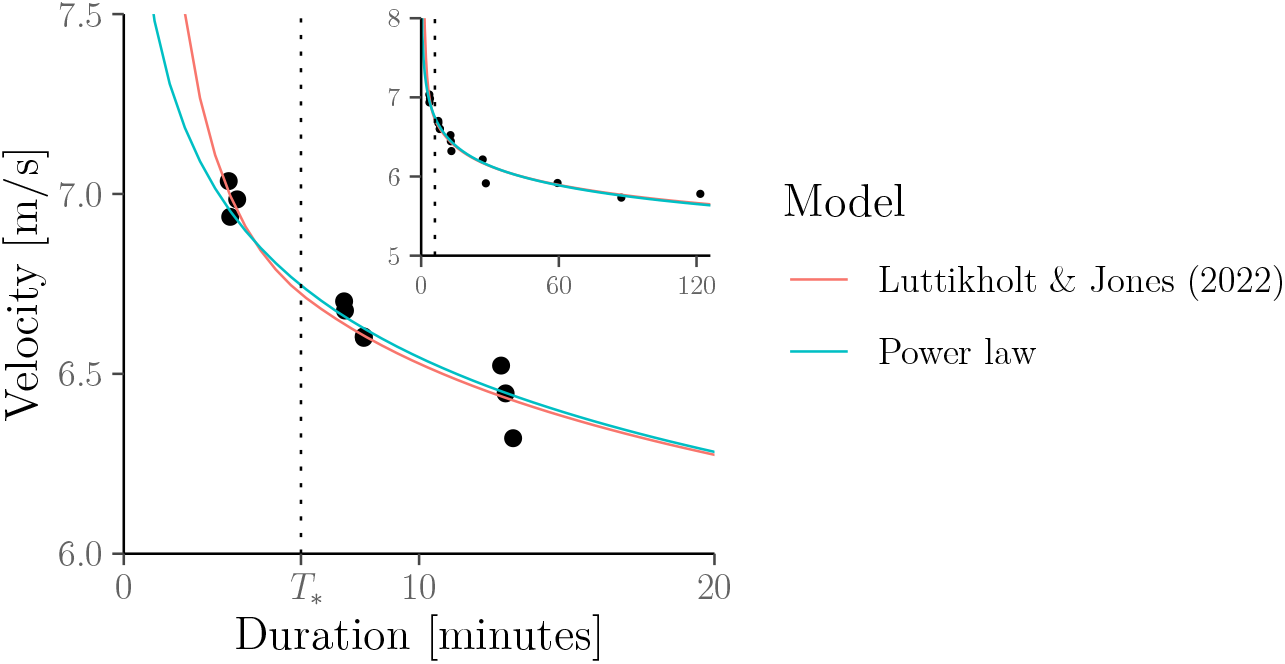
The velocity–duration relationship posited by the model from Luttikholt and Jones (2022) fitted to Eliud Kipchoge’s personal records.

## C Additional details for Section 4

### C.1 Impossibility of overpacing under the hyperbolic model

#### Proof (of Proposition 1).

Since Strategy **Pace_con_** leads to exhaustion at time *T* at which point work *W* has been accumulated, we must have

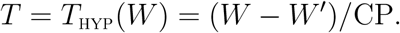

Likewise, since Strategy **Pace_var_** leads to exhaustion at time *T*_1_ + *T*_2_ at which point work *W* has been accumulated, we must have the identities:

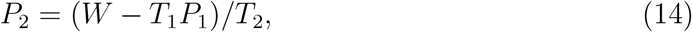

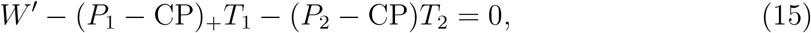

where *x*_+_ ≔ max{*x*, 0} and where we recall that *T*_1_ = *W*_1_/*P*_1_. Substituting (14) in (15) and solving for *T*_2_ then gives

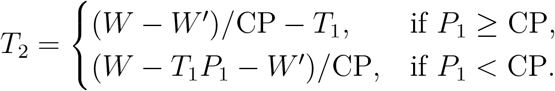

We are now ready to prove Part 1. If *P*_1_ ≥ CP, then

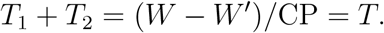

We now prove Part 2. If *P*_1_ < CP, then

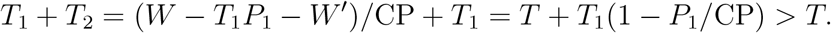

This completes the proof.

### C.2 Optimality of even pacing under the power-law model

#### Proof (of Proposition 2).

Since Strategy **Pace_con_** leads to exhaustion when work *W* has been accumulated, we must have

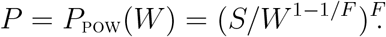

Likewise, since Strategy **Pace_var_** leads to exhaustion at time *T*_1_ + *T*_2_ at which point work *W* has been accumulated, we must have the identities:

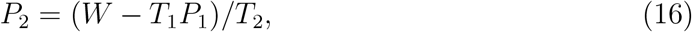

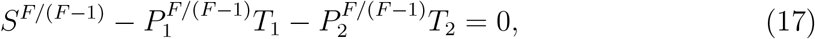

where we recall that *T*_1_ = *W*_1_/*P*_1_ and where (17) follows from our novel “rate-of-exertion” interpretation in (5) and (6) because, noting that 1/(1 – *E*) = *F*/(*F* – 1):

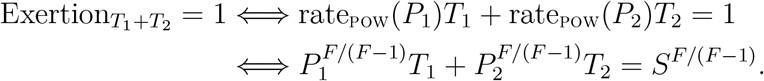

Substituting (16) in (17) and solving for *T*_2_ (now interpreted as a function of *P*_1_) then gives

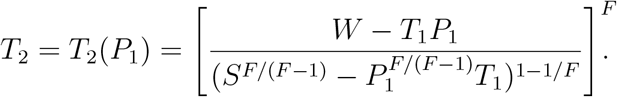

It remains to show that 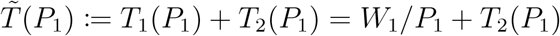 has a minimum at *P*_1_ = *P*. We have

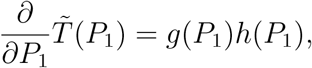

where

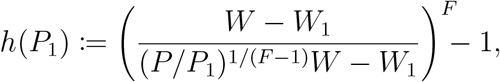

for some function *P*_1_ ↦ *g*(*P*_1_) which is strictly positive on (0, *P*_POW_(*W*_1_)) for *S* > 0, *F* > 1 and 0 < *W*_1_ < *W* (to keep the notation simple, we do not make the dependence of *g* and *h* on *W*_1_, *W*, *S* and *F* explicit in the notation). The proof is then complete since *h*((1 + *x*)*P*) is increasing in *x* and *h*(*P*) = 0; in other words, 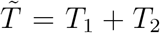 has a unique minimum at *P*_1_ = *P*. This completes the proof.

## D Additional details for Section 5

Here, we provide the mathematical details for the fatigued power–duration relationships illustrated in Figure 12. Assume that the athlete has already exercised over some duration *t* > 0 (starting from a fully rested state). Specifically, let *P*〈*s*〉 be the instantaneous power output generated by the athlete at time 0 ≤ *s* ≤ *t*. For instance, in the numerical example above, we have that *P*〈*s*〉 = *P* for any *s* > 0.

The specific form of the fatigued power–duration relationship depends on the chosen model for the fresh (i.e. non-fatigued) power–duration relationship.

- **Hyperbolic model.** If the fresh power–duration relationship is described by the hyperbolic model, and assuming for simplicity that *P*〈*s*〉 ≥ CP for all 0 ≤ *s* ≤ *t*, then the fatigued power–duration relationship is again a hyperbolic model but with *W*′ replaced by 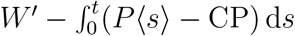.
- **Power-law model.** If the fresh power–duration relationship is described by the power-law model, the fatigued power–duration relationship is again a power-law model but with *S* replaced by 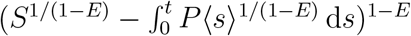.

The first result follows directly from Section 2.2.4; the second result is a consequence of the novel “rate-of-exertion interpretation” of the power-law model which we introduced in Section 2.3.4.

## E Additional details for Section 6

### E.1 Jack Daniels’ “VDOT” calculator in running

Daniels et al. (1978)^15^ performed two (non-linear) regression analyses to estimate the relationships

1. between running running velocity *V* = *D/T* [m/s] and 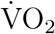 demand [ml/(kg min)]:

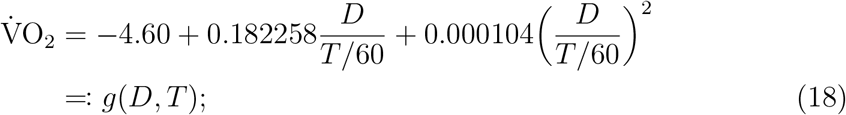
2. between race distance *D* [m] and 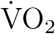 demand as a fraction of 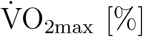 [%]:

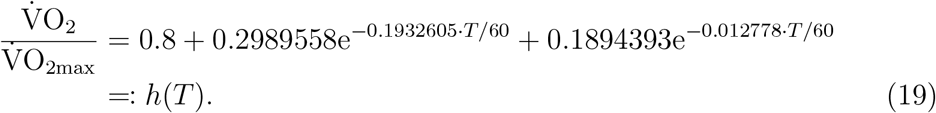

Combining (18) and (19) gives

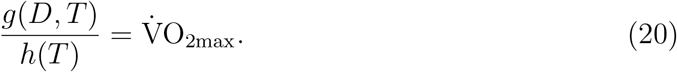

- If the athlete knows their value 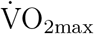, they can predict their finish time in a race over distance *D* by solving (20) for *T*. This must be done numerically using a suitable root-finding algorithm.
- If 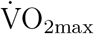 is unknown, it can be estimated from (20) using a finish time *T*_0_ for a previous race over some distance *D*_0_. That is, we estimate 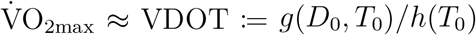. The estimate VDOT is then used place of 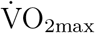 in (20) and, given some race distance *D*, we can numerically solve for *T* as before.

Solutions of (20) for various combinations of *D* and *T* can be found in the famous “VDOT” tables in Daniels (2013). These can be used if one does not wish to numerically solve (20).

As illustrated in Figure 14, except over very short durations (or distances), Daniels’ “VDOT” calculator can be viewed as an approximation the power-law model with fixed fatigue factor *F* = 1.06. This value is well within the range of values for the fatigue factor given by Riegel (1981) for running and which is also the value used by the online finish-time predictor Runner’s World Magazine (2013).

Specifically, the relationship shown in Figure 14 is derived by fixing *F* = 1.06 and then calculating the corresponding value for the parameter *S* such that the duration–distance curve of the power-law model intersects (*D*_0_, *T*_0_), where *T*_0_ ∈ {120·60, 90·60, 75·60, 60·60} [in seconds] is the time needed by a hypothetical runner to cover the half-marathon distance *D*_0_ = 21 097.5m. To calculate *S*, we use the fact that by definition of the power-law model, the velocity sustained over one mile [in m/s] is:

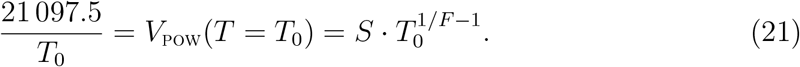

Solving (21) for *S* then gives (with *F* = 1.06):

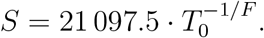

In conclusion, unless previous or predicted races are very short, the finish-time prediction *T* = *T*_0_(*D/D*_0_)^1.06^ from the power-law model (see Equation 4) is very similar to the prediction given by Daniels’ “VDOT” calculator whilst being simpler to compute because it avoids having to numerically solve (20). Though the predictions do not match exactly, any differences are likely to be negligible relative to the accuracy of estimating 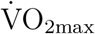 (or *S*) from a single previous race result.

### E.2 Jeff Galloway’s “Magic Mile” in running

Here, we show more formally that Jeff Galloway’s Magic Mile multipliers can be viewed as an approximation of the power-law model with *F* equal to the fatigue factor for male runners from Riegel (1981), i.e. *F* = 1.07732 (see Table 1).

Specifically, the relationship shown in Figure 15 is derived by fixing *F* = 1.07732 and then calculating the corresponding value for the parameter *S* such that the duration–distance curve of the power-law model intersects (*D*_0_, *T*_0_), where *T*_0_ ∈ {1000, 800, 600, 400} [in seconds] is the time needed by the hypothetical runner needs to cover the distance *D*_0_ = 1609.34m (one mile). To calculate *S*, we use the fact that by definition of the power-law model, the velocity sustained over one mile [in m/s] is:

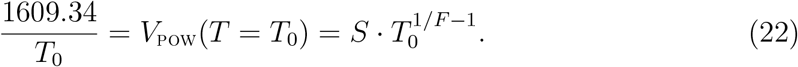

Solving (22) for *S* then gives (with *F* = 1.07732):

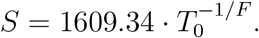

If we are just interested in the pace multipliers under the power-law model, then we do not even need to calculate *S* because these multipliers do not depend on *S*. For instance, according to the power-law model (see (10) and Section 2.1):

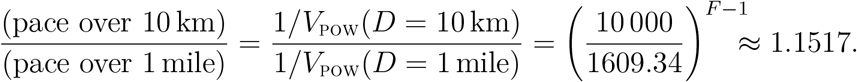

More generally, for some other distance *D* [in metres], the power-law model implies the pace multiplier (*D*/1609.34)^*F*–1^. Table 5 illustrates that this gives pace multipliers which are close to Galloway’s multipliers. The differences are likely to be negligible relative to the accuracy of estimating the mile pace.

**Table 5:** Mile-pace multiplier implied by the power-law model with *F* = 1.07732 (rounded to two digits) vs Galloway’s multiplier.

| Distance | Mile-pace multiplier |  |
| --- | --- | --- |
|  | Galloway | Power law |
| 10 km | 1.15 | 1.15 |
| 10 miles | 1.175 | 1.19 |
| Half marathon | 1.2 | 1.22 |
| Marathon | 1.3 | 1.29 |

### E.3 Kurt Jensen’s “Golden Standard” in rowing

Here, we show more formally that Kurt Jensen’s “Golden Standard” multipliers can be viewed as an approximation of the power-law model with *E* = 1/*F* = 0.85.

Specifically, the relationship shown in Figure 16 is derived by fixing *E* = 0.85 and then calculating the corresponding value for the parameter *S* such that the power–duration curve of the power-law model intersects (*T*_0_, *P*_0_), where *P*_0_ ∈ {100, 200, 300, 400} [in J/s] is the hypothetical rower can sustain over 2000m and *T*_0_ is the time needed to cover this distance. To calculate *S*, we use the fact that by definition of the power-law model:

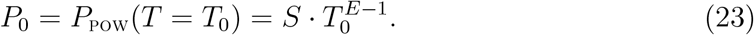

Using the conversion from (13), *T*_0_ is given by

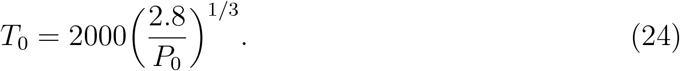

Plugging in (24) for *T*_0_ in (23) and solving for *S* then gives (with *E* = 0.85):

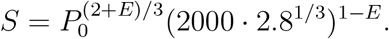

## Footnotes

1 More precisely, as discussed in the next section, this hyperbolic relationship is assumed to exist for exercise intensities in a limited range which is termed the “severe” intensity domain.

2 A power law was also suggested as an alternative to the hyperbolic model in Gorostiaga et al. (2022b) but without establishing the link to the existing literature on the power-law model and only in the context of modelling world-record performances *across* different athletes which is different from our goal of modelling the power–duration relationship of individual athletes.

3 In cycling, it has been suggested that at least the lower boundary *P*_L_ can be estimated using a three-minute all-out test (Vanhatalo et al., 2007). However this test is very arduous on athletes is known to often overestimate CP (McClave et al., 2011; Nicolò et al., 2017).

4 As we explain in Section 2.2.2 we only fit the hyperbolic model to the 2–15 minute range. Figure 17 in Appendix B.2 illustrates the compromise in fit when the model is fit to all durations.

5 The problem with the hyperbolic model is *not* that *P*_HYP_(*T*) goes to infinity as *T* goes to zero (because this is also the case for *P*_POW_(*T*)), but that *P*_HYP_(*T*) goes to infinity *too quickly*.

6 Again, as we explain in Section 2.2.2 we only fit the hyperbolic model to the 2–15 minute range. Figure 18 in Appendix B.3 illustrates the compromise in fit when the model is fit to all durations.

7 Unless it is smoothed out very heavily but then we would end up back with, essentially, a power law.

8 We stress that whilst the data in Figure 10 exhibit some kinks, e.g. at 3, 7 and 12 minutes, these should not be taken as evidence of kinks in the underlying power–duration relationship. Rather, most of the other data points likely come from sub-maximal efforts. Indeed, 3, 7 and 12 minutes are often-recommended test durations for the hyperbolic model (Karsten et al., 2015). Note also that these data were “mined” from training history so that multiple points may correspond to the same activity and then cannot all represent maximal efforts.

9 To simplify the presentation, we consider a variation in pace over only two segments. However, all results remain valid for piecewise-constant pacing strategies with more than two segments.

10 This shortcoming is even worse under a generalisation of the hyperbolic model known as the *3-parameter critical-power* model (Morton, 1996) under which an all-out pacing strategy would be *uniquely* optimal as shown in Morton (2009).

11 Fast starts were also observed to correlate with an increased time to exhaustion in a study of another seven cyclists from Jones et al. (2008). However, the meaningfulness of this finding is diminished by the fact that, although fast, slow and even starters accumulated the same amount of work over the initial two minutes, the protocol was not identical for these three conditions afterwards. That is, fast starters may have had a longer time to exhaustion simply because they continued to slow down beyond two minutes whereas the even starters had to keep their (already relatively fast) pace.

12 For instance, runners for whom this fatigue factor *F* = 1/*E* =1.06 is appropriate could estimate their 60-minute velocity as (60/20)^*E*–1^ ≈ 94% of their 20-minute velocity in analogy to to FTP.

13 For instance, runners for whom this fatigue factor *F* = 1/*E* = 1.07732 is appropriate could estimate their 60-minute velocity as (60/20)^*E*–1^ ≈ 92% of their 20-minute velocity in analogy to to FTP.

14 For instance, rowers for whom this endurance parameter *E* = 0.85 is appropriate could estimate their 60-minute power as (60/20)^*E*–1^ ≈ 85% of their 20-minute power in analogy to to FTP.

15 Equations 18 and 19 can be found in Daniels and Gilbert (1979) though we specify the durations *T* in seconds instead of minutes for consistency with the rest of our work.

